# Advanced illumination-imaging reveals photosynthesis-triggered pH, ATP and NAD redox signatures across plant cell compartments

**DOI:** 10.1101/2025.06.16.659786

**Authors:** Ke Zheng, Marlene Elsässer, Jan-Ole Niemeier, Pedro Barreto, Ana Paula Cislaghi, Minh Hoang, Elias Feitosa-Araujo, Stephan Wagner, Jonas Giese, Florian Kotnik, María del Pilar Martinez, Felix E. Buchert, José M. Ugalde, Ute Armbruster, Michael Hippler, Andreas J. Meyer, Hans-Henning Kunz, Veronica G. Maurino, Iris Finkemeier, Mareike Schallenberg-Rüdinger, Markus Schwarzländer

## Abstract

Photosynthesis provides energy and organic substrates to most life. In plants, photosynthesis dominates chloroplast physiology but represents only a fraction of the tightly interconnected metabolic network that spans the entire cell. Here, we explore how photosynthetic activity affects energy physiology within and beyond the chloroplast. We developed a new standard for the live-monitoring of subcellular energy physiology by combining confocal imaging of genetically encoded fluorescent protein biosensors with advanced on-stage illumination technology to investigate pH, MgATP^2-^ and NADH/NAD^+^ dynamics at dark-light transitions in Arabidopsis mesophyll cells. Our findings reveal a stromal alkalinization signature induced by photosynthetic proton pumping, extending to the cytosol and mitochondria as an ’alkalinization wave’. Photosynthesis leads to increased MgATP^2-^ levels in both the stroma and cytosol. Additionally, we observed reduction of the NAD pool driven by photosynthesis-derived electron export. Arabidopsis lines defective in chloroplast NADP- and mitochondrial NAD-dependent malate dehydrogenases show more reduced cytosolic NAD redox status even in darkness, highlighting the involvement of chloroplasts and mitochondria in shaping cytosolic redox metabolism via malate metabolism. Our study sets a novel methodological standard for precision live-monitoring of photosynthetic cell physiology. Applying this technology reveals signatures of photosynthetic physiology within and beyond the chloroplast with unprecedented resolution. Those signatures link photosynthetic activity and the fundamental biochemical functions of phototrophic cells.

**Significance statement:** By applying novel live microscopy monitoring using fluorescent protein biosensors in plant cells, we reveal that dark-light transitions trigger profound re-orchestration of subcellular pH, ATP and NAD redox physiology not limited to chloroplasts but extending into the cytosol and the mitochondria.

## Introduction

Photosynthesis converts sunlight into the chemical energy that sustains the vast majority of life on Earth. The light reactions of photosynthesis drive electron transport in the thylakoid membranes to deliver reducing equivalents in the form of NADPH, simultaneously establishing a proton motive force (PMF) across the thylakoid membranes via linear or cyclic electron transport that fuels stromal ATP synthesis. The photosynthetic carbon reactions of the Calvin-Benson-Bassham (CBB) cycle utilize NADPH and ATP to reduce CO2 for the synthesis of organic compounds, as building blocks and energy stores for all plant growth and survival. While electron transport activity of the photosynthetic light reactions regulates stromal metabolism through cysteine-based redox switches as mediated by the ferredoxin-thioredoxin redox system^1^ in concert with several other post-translational protein modifications^2^, stromal alkalinisation caused by proton pumping into the thylakoid lumen also directly regulates several stromal enzymes, including those involved in the CBB cycle^3–5^. Partitioning of the PMF into an electrical and chemical (proton) gradient, results in a particularly steep pH gradient across the thylakoid membrane as facilitated by its specific ion permeabilities. This is different from the mitochondrial PMF where the electrical gradient typically dominates^6^. Therefore, acidification of the thylakoid lumen and alkalinisation of the chloroplast stroma represent particularly direct changes in subcellular physiology that are intimately connected with photosynthetic function and regulation^5^. In addition to the changes in stromal cysteine-based redox status and pH as a result of photosynthetic activity, stromal ATP and NADP redox dynamics represent yet another layer with a critical impact on central metabolism. Control of metabolic energy and redox balance in the chloroplast stroma is crucial to maintain efficient carbon assimilation and to avoid photoinhibition and photodamage. The CBB cycle requires a ATP: NADPH stoichiometry of 1.5, which is not met by linear electron flow (ATP: NADPH < 1.3)^7^. Various mechanisms have evolved to adjust the ATP: NADPH ratio, by adding ATP or by dampening NADPH, or both. Those mechanisms include alternative electron flow pathways such as cyclic electron flow (CEF)^8,9^, the water- water cycle^10,11^, as well as export pathways for reducing equivalents like the triose phosphate/3-phosphoglycerate (TP/3-PGA) shuttle and the malate valve^12,13^. Even though advanced metabolic models have predicted the central importance of these pathways to maintain efficient central metabolism, validating these models with *in vivo* measurements has remained challenging^14–16^.

Historically, various methodologies have been employed to investigate subcellular pH dynamics and photosynthetic metabolism. pH-sensitive microelectrodes and chemical fluorescent dyes were early tools used for measurements in large algal and plant cells^17–22^. Yet, the difficulty in discriminating unambiguously between subcellular compartments has hampered more systematic exploration. Recently, a genetically-encoded biosensor has been established to monitor photosynthesis-associated pH dynamics in the cytosol of the cyanobacterium *Synechocystis* sp. PCC6803^23^. The luminescence of the sensor circumvents the need for fluorescence excitation but requires the loading of cells with a chemical substrate and has not yet been tested in plants. Spectroscopic methods utilizing the fluorescence and absorbance of endogenous photosynthetic pigments, like chlorophyll and carotenoids, have greatly contributed to understanding photosynthetic activity and regulation^24^. While powerful for *in vivo* measurements of photosynthetic efficiency and membrane potential, these methods are limited by challenges in interpreting data due to suboptimal properties of the pigments to act as specific sensors and the applicability to chloroplasts only, while other cell compartments remain inaccessible ^25–27^. Specialized techniques, like fast organelle fractionation and non-aqueous fractionation (NAF)^28–36^, have provided insights into bioenergetic cofactor pool dynamics, such as those of stromal ATP^31,36,37^ and NAD(P)^36,38^. Those analyses have been key for our understanding of the subcellular changes in those central cofactor pools even though the time resolution is limited.

Fluorescent protein-based biosensing offers a powerful alternative for live-monitoring dynamic changes in metabolite concentrations and physiological parameters *in planta*^39^, with subcellular resolution^40–44^. Several biosensors to monitor bioenergetic parameters related to photosynthesis, including pH^45–47^ and ATP^48–51^, are available in plants, along with sensors for NAD(P)H/NAD(P)^+52–55^ and NADPH^53,54^. Recently, also live biosensing strategies for revealing light-dependent changes in pH, ATP and NAD(P) redox status in the chloroplast and the cytosol were introduced^51,52,54^. However, capturing light-dependent dynamics has hitherto been constrained, by the difficulty of monitoring sensor fluorescence while simultaneously illuminating the tissue. First insights into light-induced dynamic changes at the subcellular level were achieved by switching between illumination and fluorescence monitoring, albeit with significant delays in the order of several seconds^50,51,54^. That temporal resolution has made it difficult to resolve reliable signatures, limiting the power of the approach to extract mechanistic insight from live responses to the illumination status of photosynthetic tissues.

Here, we aimed at unravelling the impact of photosynthetic activity on the dynamics of energy physiology in the cytosol and mitochondria. These cell compartments are separated from the chloroplasts but linked through the cellular metabolic network. We establish and optimize a novel on-stage illumination technology synchronized with confocal imaging to trigger dark-light transitions and resolve their impact on subcellular energy physiology at high flexibility and precision. By monitoring pH, MgATP^2-^ and NADH/NAD^+^ dynamics at unprecedented resolution using fluorescent protein biosensors in leaf mesophyll cells, we explore the *in vivo* impact of photosynthesis on the physiology of other cell compartments.

## Results

### On-stage illumination platform to monitor photosynthesis-related pH dynamics in the chloroplast stroma

First, we aimed at capturing the *in vivo* pH dynamics in the chloroplast stroma during dark-light transitions. As a pH sensor, we made use of a previously characterized circularly permuted Yellow Fluorescent Protein (cpYFP)^46,56,57^. We manually illuminated a cotyledon of a 5-day-old seedling mounted on the confocal microscope stage for 90 seconds (s) using white LED light at an intensity of 60 µmol m^-2^ s^-1^. This allowed tracking of the biosensor response in the mesophyll both before and after illumination. Fluorescence ratios of a chloroplast stroma-localized cpYFP (**Supplemental Figure 1a**) were increased post-illumination, indicating alkalinization of the stroma, followed by a rapid recovery in a sub-minute range (**Figure 1a**). A similar stromal alkalinization response to illumination had been shown previously by other methods^22,58,59^. However, due to the limitations associated with simultaneous quantitative fluorescence measurements of biosensors under white illumination, obtaining accurate and reliable stromal pH dynamics during illumination proved challenging. To capture the detailed pH dynamics during illumination, we established an automated on-stage illumination system synchronized with the confocal laser-scanning microscope (**Figure 1b**; for technical details see **Supplemental Table 1, Supplemental Figure 2** and **Materials & Methods**). This system allows controlled illumination at a flexible intensity that can be precisely interrupted for a short period required to record an image frame (here 1.26 s), ensuring minimal delay and a high degree of standardization (**Figure 1c**). Using this advanced setup allowed us to overcome the limitations of manual illumination (**Figure 1a**) to capture complete pH dynamics of the chloroplast stroma also in the light phase, i.e. throughout an entire cycle of dark-light-dark transition (10 minutes illumination at 60 µmol m^-2^ s^-1^; **Figure 1d; Supplemental Figure 3, 4; Supplemental Video 1**). Although interrupting illumination for 1.26 s may appear long considering the very fast biophysical processes that drive the light reactions^60^, the interruption has no obvious impact on the detected pH dynamics measured by cpYFP readout, as tested by comparison with continuous photosynthetic excitation using red LED illumination (**Supplemental Figure 5**). Those data validate the power of the new technology by providing a full picture of chloroplast-specific pH dynamics, where the stroma alkalizes at the transition from darkness to light within the first 15 s (the first data point at the illumination phase) and acidifies upon return to the darkness within 15 s (the first data point after illumination) (**Figure 1d**). An early prototype of the illumination system was made accessible to the community through a preprint to enable early adoption^61^. This prototype did not yet feature key functions, such as adjustability of light intensity, and was further improved and optimized as reported here. That said, prior versions of the technology have already enabled important new insights by the research community^62–65^.

**Figure 1.**
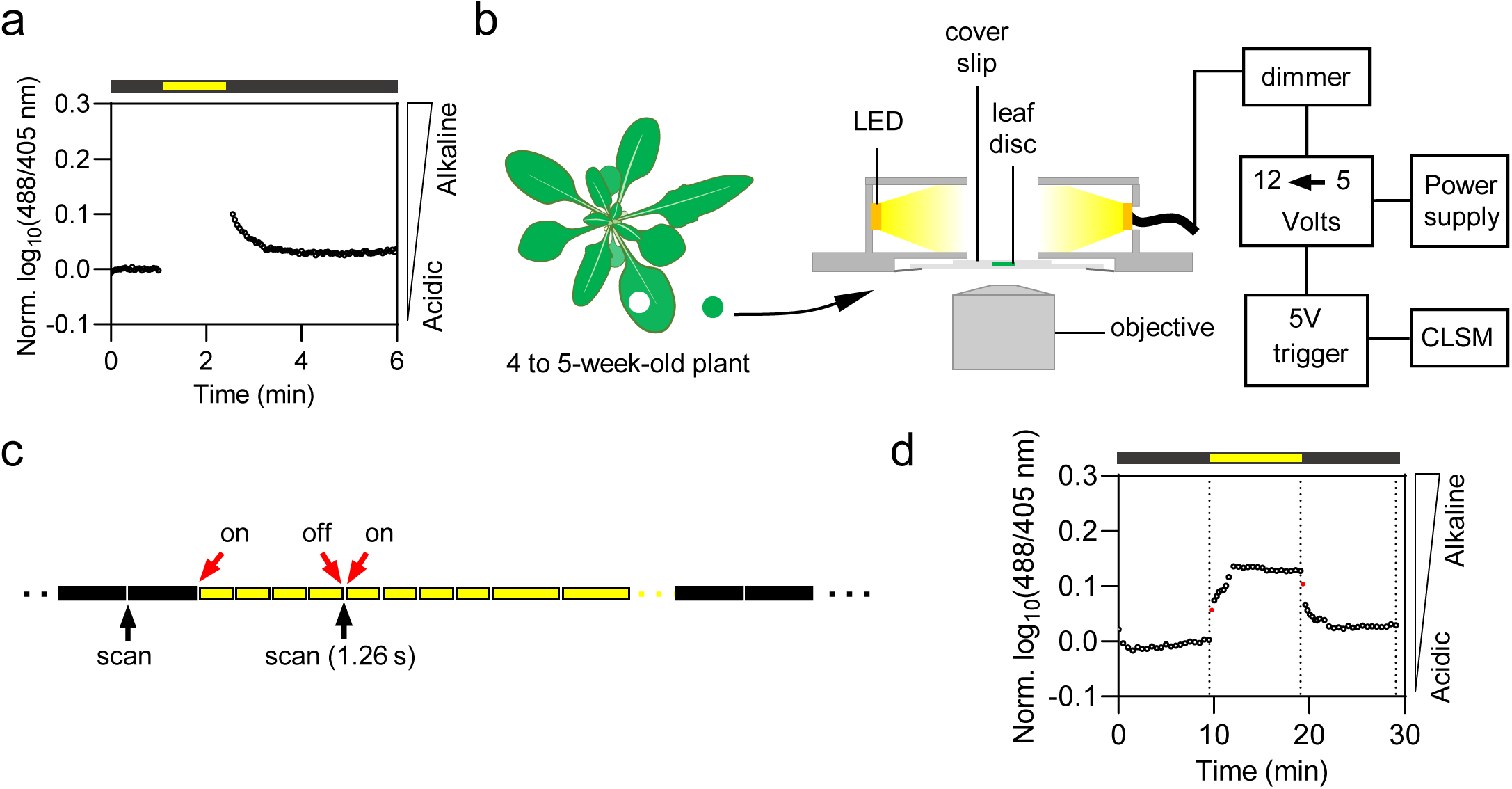
Illumination-triggered alkalinisation in chloroplast stroma monitored using the pH sensor cpYFP in *Arabidopsis thaliana* and on-stage illumination platform for confocal laser scanning microscopy. **a** Cotyledons of Arabidopsis seedlings expressing the pH-sensitive biosensor cpYFP in the chloroplast stroma were exposed to 90 seconds (s) white light (60 µmol m^-2^ s^-1^) as indicated by the yellow bar, during which no images could be captured. **b** Schematic illustration of the on-stage illumination system showing the lateral view. The aluminium chamber is equipped with a cold white LED stripe illuminating the centrally placed 4 to 5-week-old Arabidopsis leaf discs. The sample is mounted between two coverslips and immersed in perfluorodecalin to avoid hypoxia. The slide is fastened at the bottom side of the chamber, which is compatible with an inverted microscope. The 5 V trigger, controlled by the microscopy software, is converted to a 12 V signal, acting as light ON and OFF switch. The dimmer is used to adjust the light intensity from 0 to 600 µmol m^-2^ s^-1^. **c** Pseudo-continuous light created by the interruption of the illumination period for 1.26 s to scan one full frame. Illumination is controlled by the 5 V trigger signal programmed in the microscopy software. The dark phase is indicated by black bars, the light phase is indicated by yellow bars. The short bars indicate 15 s duration, the long bars indicate 30 s duration. The blank space between bars indicates a single scan event. **d** Alkalinisation in chloroplast stroma of 4 to 5-week-old Arabidopsis leaf discs in response to 60 µmol m^-2^ s^-1^ light by using the on-stage illumination system. Dash lines indicate the last data point of the dark phase and light phase. Red points indicate the first data point during and after illumination. n=3. Data are normalized to the average of the last five data points prior to illumination, log10-transformed and averaged. CLSM: The confocal laser scanning microscope

### Light intensity determines the signature of stromal alkalinization

First, we made use of the illumination system to address the remaining gap in our understanding of how varying light intensities affect stromal pH in vivo. To that end, we exposed fully expanded leaf discs of 4 to 5-week-old *Arabidopsis thaliana* plants to a 10 min period of illumination at 5, 20, 60, 200, 600 µmol m^-2^ s^-1^ (**Figure 2**). Adjustment of the optical slice along the z-axis by confocal imaging allowed the capturing of mesophyll cells below the intact epidermal cell layer. Upon illumination, the chloroplast stroma exhibited alkalinization (**Figure 2a, b**). However, during illumination, stromal pH displayed a multiphasic signature, which differed markedly depending on the light intensity (**Figure 2c**). Illumination with 5, 20 and 60 µmol m^-2^ s^-1^ triggered a rapid alkalinisation that, after two to four minutes, reached a steady-state (**Figure 2b, c**). Higher light intensities did not further increase this steady state suggesting that the steady-state stromal pH is not affected by light intensity above a certain threshold. However, the highest tested light intensity (600 µmol m^-2^ s^-1^) triggered a sharp alkalinisation spike after about two minutes of illumination (**Figure 2b; Supplemental Figure 6; Supplemental Video 2**). A similarly drastic alkalinisation spike of the chloroplast stroma was also recently observed in isolated chloroplasts at dark-light transition using fluorescent dyes^66^. Furthermore, cpYFP revealed a similar spike at a transition from low to high light ^62^. Upon transitioning from light to dark, the stromal pH rapidly dropped. While baseline was adopted by the samples illuminated with low light (5, 20 µmol m^-2^ s^-1^) at higher light fluxes (60, 200, 600 µmol m^-2^ s^-1^), the pH appeared to remain slightly elevated within the measurement window (10 min post illumination). Yet the pH elevation was variable between samples and not statistically significant (**Figure 2d**). For the measurement at 600 µmol m^-2^ s^-1^, we observed re-alkalisation after the initial acidification (**Figure 2b**). The primary driver of these pH dynamics was photosynthetic proton pumping, as evidenced by the lack of response without illumination (‘no light’; **Figure 2e**) and by the diminished pH dynamics in the presence of 3-(3,4-dichlorophenyl)-1,1-dimethylurea (DCMU), an inhibitor of photosynthetic linear electron transfer (**Figure 2f**). Saturated inhibition of photosynthesis by DCMU in the leaf samples was validated by measurement of reduced quantum yields of photosystem II and depleted energy-dependent non-photochemical quenching (**Supplemental Figure 7**).

**Figure 2.**
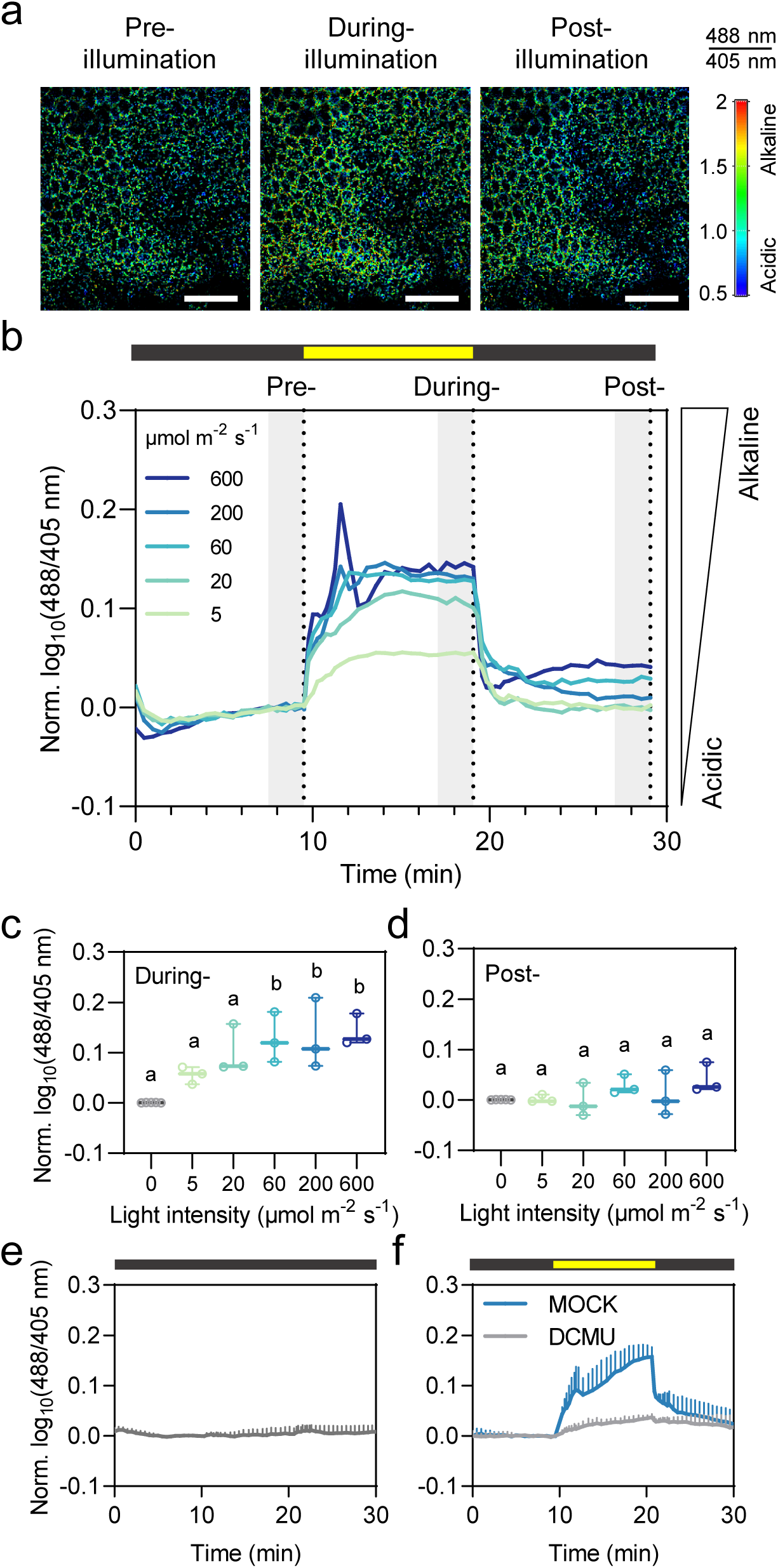
Light intensity-dependent alkalinisation in chloroplast stroma. **a** Ratiometric images of mesophyll expressing cpYFP in the chloroplast stroma taken before, during and after illumination at 600 µmol m^-2^ s^-1^ (data points are displayed as dash lines in **b**). Scale bars: 200 µm. 488/405 nm excitation ratio at 508-535 nm emission is indicated in false color. **b** Alkalinisation in chloroplast stroma in response to different light intensities (indicated by yellow bars) for 10 min. n = 3. The grey patches represent the last five data points from each phase, which is used in the following analysis. **c** The pH of the chloroplast stroma increases at the end of illumination, correlating positively with light intensities. **d** Slightly elevated stroma pH after illumination. **e** ‘No light’ control in the chloroplast stroma-located cpYFP. n = 4. **f** 3-(3,4-dichlorophenyl)-1,1-dimethylurea (DCMU) control and the respective ‘MOCK’ treatment in the chloroplast stroma-located cpYFP. Leaf discs were pre-incubated in 20 µM DCMU and 0.2% (v/v) EtOH, or 0.2% (v/v) EtOH, respectively, for 35 to 45 min. n = 3-4. Light intensity is 200 µmol m^-2^ s^-1^. Data points of 0 µmol m^-2^ s^-1^ in **(c)** and **(d)** are from ‘Pre-’ in **(b)**. The difference of sensor ratio in varying light intensities was tested by One-way ANOVA with Tukey‘s multiple comparisons. Distinct letters indicate significant differences (p < 0.05). For panels **(b)-(f)**, data are normalized to the average of the last five data points prior to illumination (‘Pre-’), log10-transformed and averaged. Error bars = SD.

### Photochemical activity triggers a stromal alkalinization signature that spreads beyond the chloroplast

While chloroplast stroma alkalinization is intrinsic to current models of photosynthesis, early reports, based on live measurements with pH-microelectrodes, e.g. in *Riccia fluitans*^17^ or fluorescent dyes, e.g. in *Pelargonium zonale*^22^, proposed that also cytosolic pH may be affected by photosynthetic activity. However, the temporal and subcellular resolution of those measurements was insufficient for further dissection. To investigate the potential impact of photosynthetic activity on subcellular pH beyond the chloroplast, we next compared cytosolic and stromal pH dynamics at 20, 200 and 600 µmol m^-2^ s^-1^ (**Figure 3a-f**). The cytosol-located pH biosensor cpYFP (**Supplemental Figure 1b**), indicated cytosolic pH signatures that were similar to those of the stroma only at 20 µmol m^-2^ s^-1^ (**Figure 3d**). Higher light intensities induced an alkalinization spike at around 1 min after the start of illumination. Then pH dropped to pre-illumination levels (**Figure 3e, Supplemental Figure 8a-c, Supplemental Video 3**). This transient pH event is in line with early observations using microelectrodes in algal and moss cells^17,20,67^. Similar observations were made with cytosolic pH-GFP, which was used as a distinct pH sensor with a lower p*K*a^68^ (**Supplemental Figure 9, Supplemental Video 4**). pH decline after the initial alkalinization spike also occurred under lower light intensity (60 µmol m^-2^ s^-1^), albeit to a lesser extent (**Supplemental Figure 10a-c**). Following the initial transient alkalinization event, cytosolic pH increased gradually for the duration of the illumination phase (**Figure 3e, f; Supplemental Figures 8a-c; 9**). Upon the transition from light to dark, the cytosol acidified sharply, followed by a return to similar pH values as towards the end of the illumination phase. Maintenance of a more alkaline cytosolic pH within the first minutes after illumination(**Figure 3e, f**) is in agreement with early microelectrode analyses observations^17,20,67^. Given the tight association of chloroplasts and mitochondria^69,70^ as well as the connection between both the chloroplast stroma and the mitochondrial matrix with the cytosol via a large set of membrane transport proteins, we hypothesized that also mitochondrial pH may be influenced by photosynthetic activity. Monitoring pH dynamics of the mitochondrial matrix *in vivo* had remained impossible via microelectrodes due to the small organelle size. Remarkably, mitochondrial pH, as monitored by a matrix-localized cpYFP pH biosensor (**Supplemental Figure 1c**)^46^, showed rapid alkalinization in synchrony with the chloroplast stroma and cytosol at onset of illumination (**Figure 3g-i, Supplemental Figure 8d-f, Supplemental Figure 10d-f, Supplemental Video 5**). Even though a slight transient acidification was apparent directly after the initial alkalinisation spike, pH increased throughout the illumination period, and dropped upon the light-to-dark transition. As such, mitochondrial matrix pH resembled a mix of the signatures observed in the chloroplast stroma and the cytosol. The pH signatures in both the cytosol and mitochondria were absent in dark-control samples and diminished by DCMU (**Supplemental Figure 11**). Next, we sought to understand the characteristics of the different subcellular pH signatures and analysed their individual properties in a comparative manner (**Figure 3j-o**). Notably, the chloroplast stroma exhibited the fastest alkalinization among the three compartments at all light intensities (**Figure 3j- l**), in agreement with the likely role of the thylakoids as a primary proton sink. The slight delay in the alkalinization of the mitochondria and the cytosol plausibly reflects several processes including proton exchange across membranes^5^, as well as distinct pH buffering and resting pH of different cell compartments^71^. During light exposure, all compartments showed higher excitation ratios, indicating more alkaline conditions than before illumination (**Figure 3m-o**). Additionally, after illumination, all compartments also showed noticeably higher pH compared to the pre-illumination phase, particularly at the highest light flux at 600 µmol m^-2^ s^-1^ (**Figure 3o**). These observations showcase the power of the on-stage illumination technology in detecting pH dynamics across different cell compartments. They reveal a global impact of light-dependent photochemical activity on cellular pH dynamics, including those of the chloroplast stroma but also of the cytosol and the mitochondrial matrix, that had hitherto been impossible to resolve.

**Figure 3.**
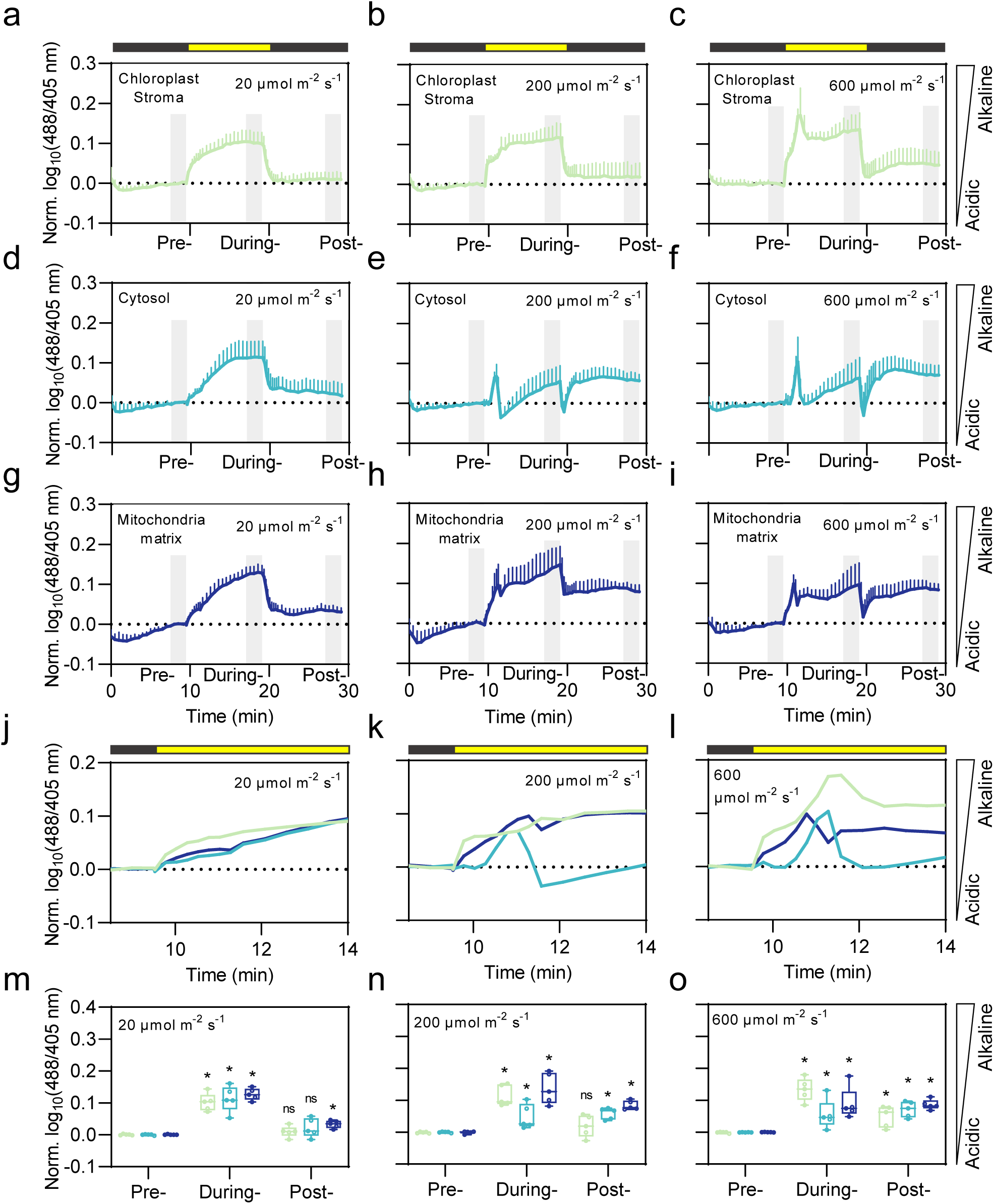
Light intensity-dependent alkalinisation and acidification in chloroplast stroma, cytosol and mitochondrial matrix. **a-c** pH in chloroplast stroma (green), **d-f** cytosol (cyan) and **g-i** mitochondria (blue) in response to 20, 200 and 600 µmol m^-2^ s^-1^ light (indicated by yellow bars) for 10 min. n = 5. **j-l** Magnification of dark-to-light transition of three compartments. **m-o** Comparison of the Pre-, During and Post-illumination phases (defined as the mean of the last five measurement points of each phase, indicated by grey shading). Two-way ANOVA with Dunnett‘s multiple comparisons test to compare During- and Post-illumination of each line to Pre-illumination. ns, not significant; * p < 0.05. Data are normalized to the average of last five data points prior illumination, log10-transformed and averaged. Error bars = SD. The dotted horizontal lines indicate log10(488/405 nm) = 0.

### MgATP^2-^ concentrations increase during active photosynthesis in the chloroplast stroma and the cytosol

The linkage of cytosolic and mitochondrial pH to photosynthesis-induced changes in stromal pH prompted us to extend our analysis to ATP, because ATP production and consumption are intimately linked with pH gradients across cellular membrane systems through ATP-synthases/H^+^-ATPases^72,73^. Mitochondrial respiration is the primary source of cytosolic ATP^16,74^, and there has been no evidence of direct ATP export from the chloroplast stroma in mature leaves^50,75–77^. To assess stromal and cytosolic ATP dynamics in response to changes in photosynthetic activity, we took advantage of Arabidopsis plants expressing the Förster Resonance Energy Transfer (FRET) sensor ATeam1.03-nD/nA which responds specifically to MgATP^2-^, as the major physiologically relevant form of ATP in cells ^48,78,79^ (**Supplemental Figure 12**). The standardized 10 min illumination regime used for monitoring pH dynamics revealed an increase in FRET ratio in the stroma indicating a rise in stromal MgATP^2-^ concentration reaching a first maximum after approximately 1 min (**Figure 4a, b; Supplemental Figure 13, Supplemental Video 6**). The apparent dynamics of stromal MgATP^2-^ were similar under all tested light intensities (**Figure 4b**). Even upon the lowest illumination intensity (5 µmol m^-2^ s^-1^) the initial increase occurred; yet the subsequent gradual increase was only observed at the higher intensities. The absence of any response in the ‘no light’ and DCMU controls demonstrates that the shift towards higher levels in stromal ATP is caused by photosynthetic electron transport (**Figure 4c, d**). Since both ATP production by the light reactions and ATP consumption by the CBB cycle are activated by illumination, the observed ATP dynamics validates that the stroma operates at increased adenylate charge in the light *in vivo.* Increased adenylate charge is in line with previous indirect estimations via spectrophotometric-based analyses^80^ and analytical measurements following cell fractionation^28,31,34,37,81^. The apparent increase in MgATP^2-^ concentration is likely to represent a pronounced shift in adenylate charge considering that MgATP^2-^ concentrations are buffered by stromal adenylate kinase activity^82,83^. Interestingly, cytosolic MgATP^2-^ levels also increase in the light, which we validated through illumination at 60 µmol m^-2^ s^-1^ (**Figure 4e, Supplemental Video 7**). Yet, the light-to-dark transition did only cause a short-lived decrease in cytosolic MgATP^2-^ followed by a further increase after 1 min. This unexpected behaviour was reproducible in all replicates and absent in the ‘no light’ and DCMU controls (**Supplemental Figure 14**). We further investigated cytosolic MgATP^2-^ levels under higher light intensities and surprisingly observed a slight decrease in FRET ratio during illumination. However, this decrease persisted in DCMU pre-incubated samples and was hence not caused by photosynthetic activity (**Supplemental Figure 15a, b**). A plausible explanation is provided by the known temperature sensitivity of ATeam biosensors^78^ (increasing temperature causes an increase in *K*d). The slight decrease in sensor FRET-ratio may be attributed to a rise in tissue temperature during illumination. Indeed, under the highest light flux (600 µmol m^-2^ s^-1^), we measured an increase in temperature in the location of the sample within the on-stage illumination system by up to 6 degrees (**Supplemental Figure 15c**). Similarly, increasing temperature in the dark also caused a decrease in sensor ratio (**Supplemental Figure 15d**). The temperature effect on ATeam could also influence the stromal MgATP^2-^ measurements, where it would lead to an underestimation of the increase in ATP. FRET values of the non-normalized log10(Venus/CFP) data from 0.4 to 0.48 in the cytosol and 0.18 to 0.2 in the stroma indicate generally lower MgATP^2-^ concentrations in the stroma than in the cytosol (**Figure 4f**). Distinct MgATP^2-^ pool sizes between the stroma and the cytosol in living mature leaves are in line with recent estimates of about 0.2 mM of stromal MgATP^2-^ in the dark and 0.5 mM in the light, and about 2 mM of cytosolic MgATP^2-36,48,50^.

**Figure 4.**
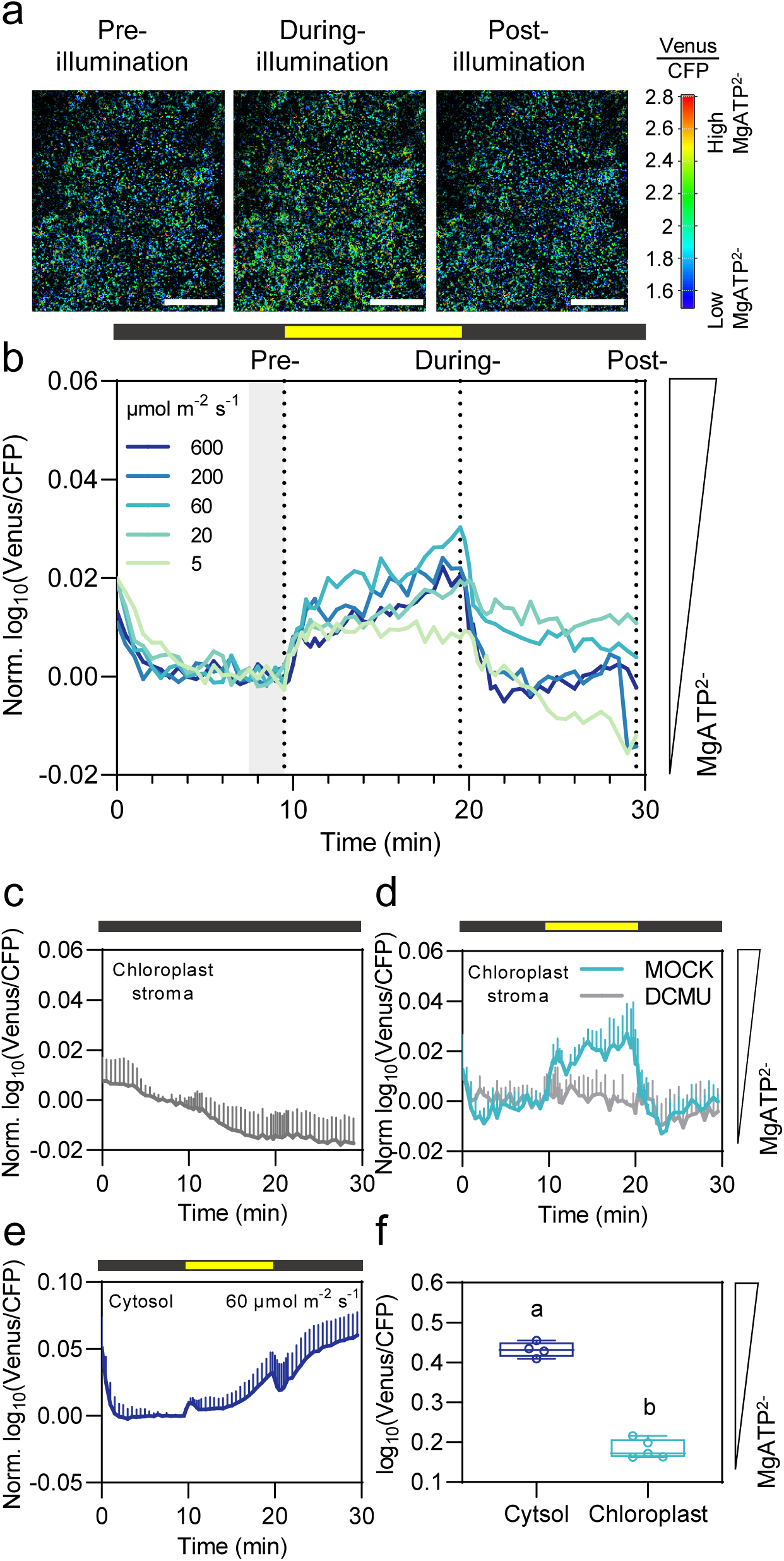
Chloroplast stromal and cytosolic MgATP^2-^ dynamic in response to dark-light transitions. **a** Ratiometric images of the final image taken before, during and after illumination at 60 µmol m^-2^ s^-1^ of mesophyll expressing ATeam1.03-nD/nA in the chloroplast stroma (data points are displayed as dash lines in **b**). Scale bars: 200 µm. The ratio of Venus/CFP is indicated in false color (excitation: 445 nm, Venus emission: 526-561 nm; CFP emission: 464-499 nm). **b** MgATP^2-^ increase in chloroplast stroma in response different light intensities (indicated by yellow bars) for 10 min. The grey patches represent the final five data points from two dark phases, which is used in the following analysis. n = 3. **c** ‘No light’ controls in the chloroplast stroma-located ATeam1.03-nD/nA. n = 5. **d** ‘DCMU’ control and the respective ‘MOCK’ treatment in the chloroplast stroma-located ATeam1.03-nD/nA. Leaf discs were pre-incubated in 20 µM DCMU and 0.2% (v/v) EtOH, or 0.2% (v/v) EtOH, respectively, for 35 to 45 min. n = 3. Light intensity is 60 µmol m^-2^ s^-1^. **e** Cytosolic MgATP^2-^ dynamics in response to 60 µmol m^-2^ s^-1^ light (indicated by yellow bars) for 10 min. n = 4. Data are normalized to the average of last five data points prior illumination, log10-transformed and averaged. Error bars = SD. **f** Comparison of unnormalized chloroplast stroma and cytosolic sensor ratio in Pre- and Post- illumination (indicated in **b**). Light intensity is 60 µmol m^-2^ s^-1^. Two-way ANOVA with Tukey‘s multiple comparisons test to compare sensor ratio difference between chloroplast stroma and cytosol. Distinct letters indicate significant differences (p < 0.05).

### A photosynthesis-dependent redox signature of the cytosolic NAD pool

We next aimed to assess the subcellular dynamics of metabolic redox equivalents as a key output of photosynthetic activity. While several redox shuttle systems, such as the malate valve, have been studied intensely^12,13^, assessing their real-time *in situ* impact has been difficult to dissect. To monitor the export of reducing equivalents into the cytosol, we focussed on the redox dynamics of the cytosolic NAD pool using the NADH/NAD^+^ biosensor Peredox-mCherry (**Supplemental Figure 16**)^52,84^. While previous studies have noted the reduction of the cytosolic NAD pool following illumination, the exact dynamics of the NAD redox status during illumination have yet to be elucidated^52^. Employing the advanced illumination system, we exposed Arabidopsis leaf discs expressing cytosolic Peredox-mCherry to a 15 min period of light (initial tests using 10 min revealed the need for longer light exposure to reach a steady-state). As a result, we observed characteristic signatures, indicating pronounced NADH/NAD^+^ redox dynamics (**Figure 5**). Rapid reduction of the cytosolic NAD pool was evident from the immediate increase in biosensor ratio upon illumination, which signifies the rapid export of photosynthesis-derived reducing equivalents from the chloroplast stroma (**Figure 5a, b; Supplemental Figure 17, 18; Supplemental Video 8**). The initial reduction that was similar in rate between 20 and 600 µmol m^-2^ s^-1^ was followed by a transient oxidation after about 30 – 60 s of illumination, which was reached faster at increasing light intensities (**Figure 5b, c**).

**Figure 5.**
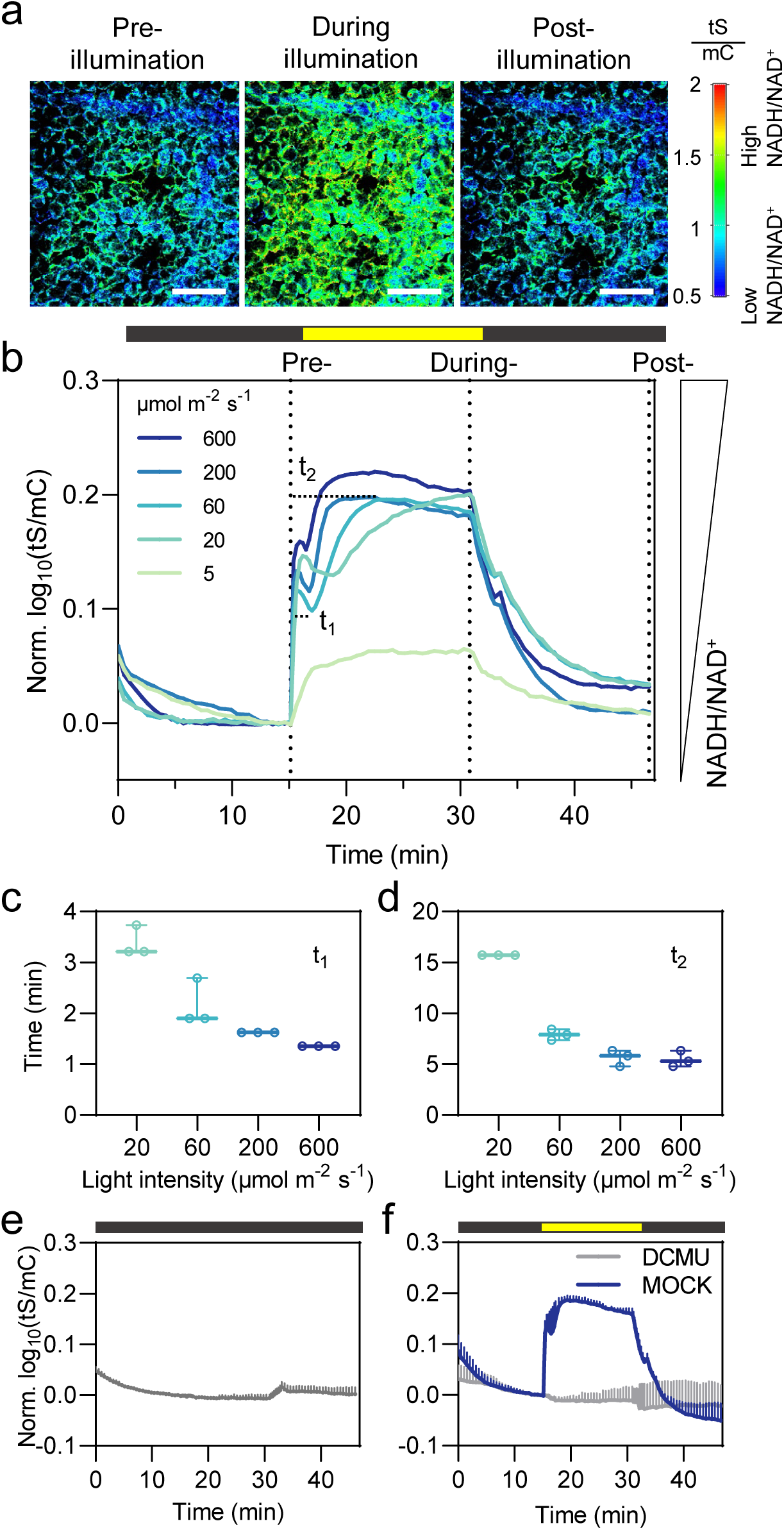
Light intensity-dependent NAD pool reduction in the cytosol. **a** Ratiometric images of the final image taken before, during and after illumination at 20 µmol m^-2^ s^-1^ of mesophyll cells expressing Peredox-mCherry in the cytosol (data points are displayed as dash lines in **b**). Scale bars: 200 µm. The ratio of tSapphire/mCherry (tS/mC) is indicated in false color (tS: excitation: 405 nm, emission: 499-544 nm; mC: excitation: 561 nm, emission: 588-624 nm). **b** NAD reduction in the cytosol in response to different light intensities (indicated by yellow bars) for 15 min. 5 and 600 µmol m^-2^ s^-1^ n = 2, other light intensities n=3. t1 and t2 indicate the duration to reach the minimum and plateau during illumination separately in the case of 60 µmol m^-2^ s^-1^. **c** The time point (t1) of tS/mC ratio to reach its minimum under different light intensities. **d** The time point (t2) of tS/mC ratio to reach plateau under different light intensities **e** ‘No light’ control in the cytosol-located Peredox-mCherry leaves n = 4. **f** ‘DCMU’ control and the respective ‘MOCK’ treatment in the cytosol-located Peredox- mCherry leaves. Leaf discs were pre-incubated in 20 µM DCMU and 0.2% (v/v) EtOH, or 0.2% (v/v) EtOH, respectively, for 35 to 45 min. Light intensity is 600 µmol m^-2^ s^-1^. N = 3. For panels **(b)-(f)**, data are normalized to the average of last five data points prior illumination, log10-transformed and averaged. Error bars = SD. Red.: Reduction; Ox.: Oxidation.

Subsequently, reduction increased gradually at rates proportional to intensity (**Figure 5b, d**), and maximum reduction was reached from approximately 5-10 min of light exposure on. Only the lowest light intensity (5 µmol m^-2^ s^-1^) showed a slower initial reduction and a lower amplitude that lacked any distinction of the two reduction phases. After illumination, cytosolic NAD re-oxidized and returned to a status similar to pre-illumination. The NAD redox signatures did not occur in the ‘no light’ and the DCMU controls (**Figure 5e, f**), demonstrating that cytosolic NAD reduction was caused by photosynthetic electron transfer activity.

### Loss of organellar malate dehydrogenases affects cytosolic redox balance

To pinpoint the mechanisms underpinning the cytosolic NAD redox signatures, we focussed on the malate valves as a candidate mechanism to accommodate major flux of reducing equivalents between the chloroplast stroma or the mitochondrial matrix and the cytosol^13^. Considering photorespiratory glycine oxidation and the resulting production of reductive power in the form of NADH in the mitochondrial matrix under illumination, the observed cytosolic NAD reduction signature (**Figure 5b**) may reflect the export of reducing equivalents from the chloroplasts, the mitochondria or both^16,54^. In such a scenario, the organellar malate valves are prime candidates to mediate the redox connectivity between the plastid stroma, the mitochondrial matrix and cytosol^13^. We, therefore, tested the significance of malate valve capacity for the reduction of the cytosolic NAD pool. We selected and validated previously established Arabidopsis loss-of-function lines of three organellar key enzymes that contribute to the malate valves, namely the mitochondrial NAD-dependent malate dehydrogenases 1 and 2 (mMDH1, At1g53240; mMDH2, At3g15020)^85^ and chloroplast NADP-dependent MDH (NADP-MDH) (At5g58330)^86^, and equipped them with the cytosolic Peredox-mCherry biosensor (**Figure 6**). Strikingly, the overall reduction signature of the cytosolic NAD pool during illumination, including its biphasic pattern, remained largely unchanged between wild type, *mmdh1*, *mmdh2* and *nadp-mdh* (**Figure 6a, b**). However, the amplitude of the NAD reduction signature was diminished in all three mutants. Inspecting the non-normalized data revealed that the sensor ratios in all three malate dehydrogenase loss-of-function lines were notably higher at baseline, indicating a more reduced cytosolic NAD pool already at resting levels, as well as the entire experimental illumination regime, when compared to the wild type (**Figure 6c-f**). The *mmdh1* background, lacking mitochondrial MDH1, displayed the smallest amplitude change over the entire period, while loss of NADP-MDH in the chloroplast stroma had the strongest impact on absolute reduction in the darkness and in the light (**Figure 6g, h**). The observation of altered cytosolic NAD redox status even in the dark, was strictly reproducible across experiments, yet surprising considering that NADP-MDH has been extensively studied as an example of a redox-regulated enzyme that is active only in the light^87^. NAD redox status recovered after illumination, returning to similar, genotype-specific values as prior to illumination (**Figure 6g, h**). In summary, our results indicate that constraining MDH activity in mitochondria and chloroplasts does not impair the general functionality of the malate valves at dark-light transitions. Rather, a higher NAD reduction load in the cytosol is pinpointed as a common phenotype to all three *mdh* backgrounds, possibly as a consequence of the constraint on organelle MDH capacity to cause re-routing of organellar malate reduction to the cytosol (**Figure 6i**).

**Figure 6.**
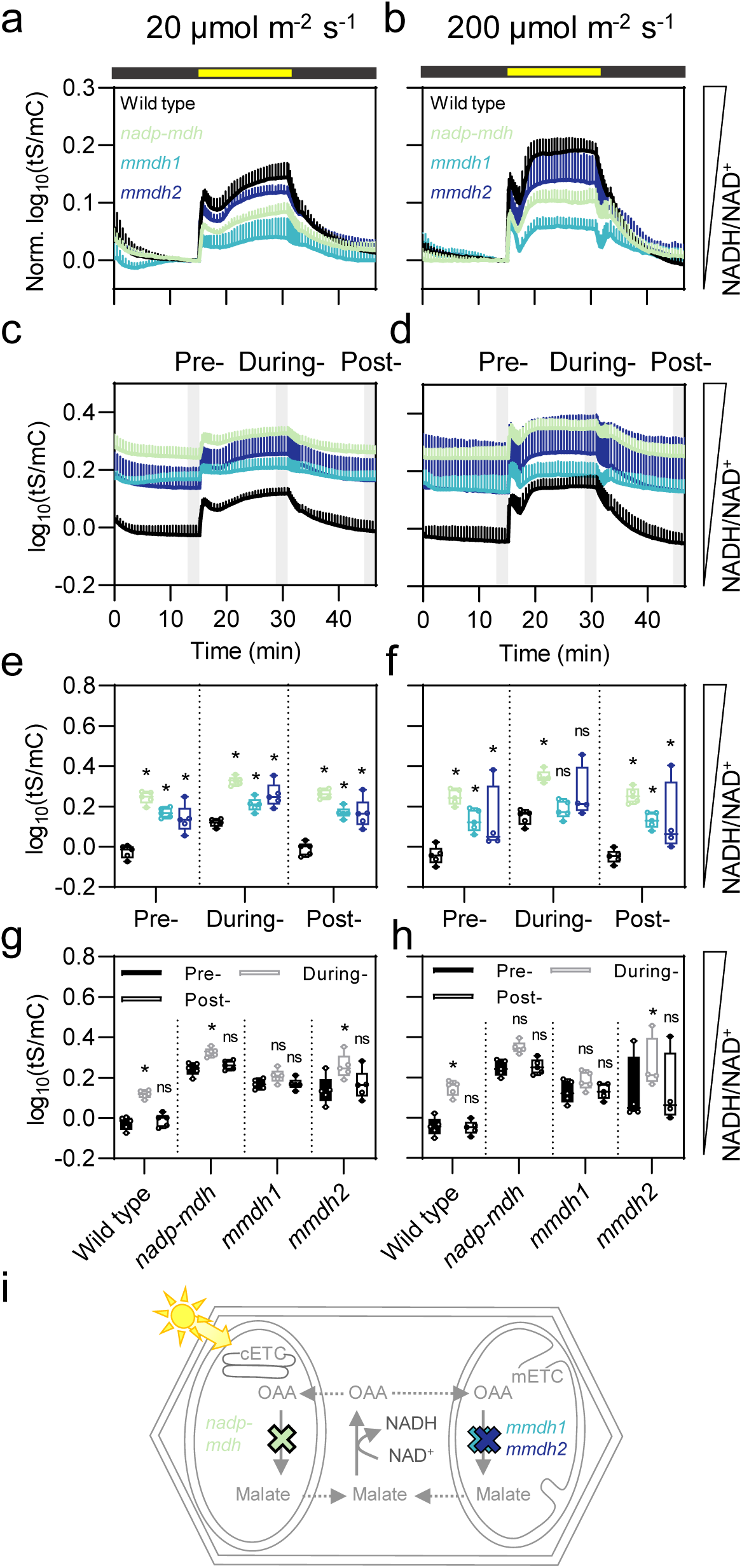
Reduction of the cytosolic NAD pool in organellar *mdh* lines in the dark- light transitions. **a, b** NAD reduction in the cytosol of the genotypes tested in response to 20 and 200 µmol m^-2^ s^-1^ light (indicated by yellow bars) for 15 min. n = 5. Data are normalized to the average of the last five data prior illumination, log10-transformed and averaged. Without normalization, data are shown in **(c)** and **(d)**. The grey patches represent the last five data points from each phase, which is used in the following analysis. **e, f** Comparison of NAD redox of *mdh* lines to wild type in the Pre-, During and Post- illumination phases (data points indicated in **c** and **d**). Two-way ANOVA with Dunnett‘s multiple comparison test. **g, h** Comparison of NAD redox in During- and Post- illumination of each line to Pre-illumination phase (data points indicated in **d** and **e**). Two-way ANOVA with Dunnett‘s multiple comparisons test. Data are log10- transformed, averaged. ns, not significant; * p < 0.05. Error bars = SD. Dash lines are used to separate different phases or genotypes. **i** Interference with the subcellular malate metabolism by knockout of the *nadp-mdh* (green), *mmdh1* (cyan) or *mmdh2* (blue). A functioning malate shuttle system leads to malate export from the organelles (dotted arrow) and conversion to oxaloacetate (OAA) in the cytosol and concomitant reduction of NAD^+^. *nadp-mdh*: *plastidial NADP-dependent malate dehydrogenase*. *mmdh1*: *mitochondrial NAD-dependent malate dehydrogenase 1*. *mmdh2*: *mitochondrial NAD-dependent malate dehydrogenase 2*. cETC: chloroplast electron transport chain. mETC: mitochondrial electron transport chain.

## Discussion

### Advanced illumination imaging setup to unlock subcellular photosynthetic physiology

Here we developed and deployed a novel illumination platform to reveal the *in vivo* pH, ATP, and NAD redox signatures across different cellular compartments that are caused by increasing degrees of photosynthetic activity in Arabidopsis leaf mesophyll. The platform offers high temporal resolution at the cell compartment level, thus revealing detailed physiological signatures that are triggered by photosynthetic activity. Their compartment-specific fingerprints offer insights into how the local biochemical machinery integrates the changes in physiological status *in vivo* that confirm results obtained by other approaches. They also provide several insights that are novel. The platform introduced here is expandable and offers several options for targeted refinements depending on the specific question investigated. The integrated light switching provides a major improvement compared to previously used systems, which suffered from seconds of lag time between illumination and recording, which can be decisive for rapidly changing parameters such as pH^50,51,54^. The image acquisition interval can be further minimized down to the millisecond range if needed by adjusting the microscope settings for faster laser scanning (**Figure 1c**). Using red instead of white light illumination even offers biosensing during active photosynthesis (**Supplemental Figure 5**), and has very recently been exploited in combination with an earlier version of the system introduced here^62,64^. By optimizing previous prototypes^61^, the advanced on-stage illumination system now includes the option of continuously adjustable light intensity (0 to 600 µmol m^2^ s^-1^) and allows flexible exchange of LEDs to support any illumination light quality. Thus, it provides a platform technology to address a broad set of questions on how light and/or photosynthesis affect subcellular physiology and metabolism. It may also be applied beyond photosynthesis research, e.g. for assessment of photoreceptor responses or optogenetics. Remaining technical limitations such as the elevated temperatures that we observed at particularly high light fluxes (**Supplemental Figure 15c**) may be addressed through further optimization of design, such as the use of high thermal conductance materials or implementation of active temperature control by heating and cooling. The illumination setup may further be combined with a perfusion option for chemical treatments or atmospheric control to control CO2 and O2 availability in order to move closer towards comprehensive environmental control live on-stage. While the setup is suitable for microscopy stages, its principles may be applied to other optical devises that are routinely used for fluorescence-based biosensing, such as plate reader-based fluorimeters which enable higher sample throughput.

### A photosynthetic ‘pH wave’ affects large parts of the cell

The advanced illumination imaging setup revealed multiphasic stromal pH signatures in live Arabidopsis mesophyll in response to photosynthetic activity. Notably, at the onset of the higher light fluxes that we assessed here (200 and 600 µmol m^-2^ s^-1^), stromal pH exhibited a sharp spike before stabilizing (**Figure 2b; Supplemental Figure 6**). A similar alkalinisation spike in the stroma was recently observed at a low-to-high light transition from 90 to 900 µmol m^-2^ s^-1^ as enabled by an earlier prototype of the illumination imaging setup^62^. Candidate mechanisms that may shape the alkalinisation spike include ion fluxes across the thylakoid membrane, but also changes in cyclic electron transport rate, energy-dependent quenching (qE) and photosynthetic control. Recently, simultaneous monitoring of fluorescent protein sensors and chlorophyll fluorescence analysis was introduced^88,89^, which may proof useful to shed light on the mechanism(s) that underpin the alkalinisation spike in the future.

Like stromal pH dynamics, cytosolic pH also exhibited significant changes under illumination despite being well-buffered at approximately 20 to 80 mM H^+^ per pH unit^90^. Light-induced alkalinization still occurs in the cytosol, indicating a significant proton efflux between the cytosol and other compartments under illumination. Illumination-induced alkalinization of the cytosol was also observed in the aquatic liverwort *Riccia fluitans*, where large cells allow precise insertion of pH-sensitive microelectrodes^17^. The signature largely resembled the properties of the alkalinization described in *Riccia fluitans*, including an amplitude of about 0.4 pH units^17^. In higher plants, indirect indications for light-induced changes in cytosolic pH were reported, such as light-induced depolarisation of the plasma membrane^91,92^. Further, indirect evidence was reported from the use of fluorescent dyes, such as pyranine or 5-carboxyfluorescein diacetate, in illuminated leaves or protoplasts^19,22^. The independent observation of cytosolic alkalinization in evolutionary distant plant systems using orthogonal approaches suggests that a cellular ‘pH wave’ may be a ubiquitous event that occurs at transitions in photochemical activity. Even though the conversion of sensor ratios obtained *in vivo* to absolute pH values should be regarded with caution since additional assumptions are introduced, calibration curves based on leaf extract^44^ and purified sensor protein^56^ allow us to estimate the observed light-induced pH shift at around 0.4 pH units in all three compartments. After the initial transient alkalinization at the onset of illumination, cytosolic pH as revealed by cpYFP (and pH-GFP as an independent pH sensor with a lower p*K*a^68^), decreased sharply reaching pH levels below the pre- illumination baseline (**Figure 3d-f; Supplemental Figure 9**). While it is evident that the pH signatures that are induced by photosynthesis are distinct between stroma and cytosol, pinpointing the exact mechanisms that (i) connect both signatures and (ii) shape the cytosolic signature is complicated by the large number of mechanisms that may affect pH. At the onset of illumination, proton-pumping across the thylakoid membrane by photosynthetic electron transport causes stromal alkalinisation, which in turn changes the pH gradient across the inner envelope. It is plausible to assume that various proton-coupled solute transporters in the inner envelope will be able to mediate proton entry from the cytosol into the stroma, leading to the initial cytosolic alkalization event. In turn, cytosolic alkalinisation is likely to lead to the import of protons from other cell compartments including from the mitochondria (**Figure 3g-i**), as well as the vacuole and the apoplast. Indeed, alkalization at the onset of illumination was previously observed in other cell compartments, such as the apoplast in *Vicia faba* leaves^93^. At light-to-dark transition, the primary cause for the acidification is the stop of photosynthetic proton pumping across the thylakoid membrane. This is followed by the re-set of cellular pH homeostasis, which is likely to involve the re-distribution of different solutes and may explain the transient acidification observed in the stroma and mitochondria post-illumination (**Figure 3c, i**). The mitochondrial matrix exhibited transient alkalinization similar to that observed in the cytosol at the onset of illumination (**Figure 3g-i**), suggesting that cytosolic pH has a direct impact on matrix pH, possibly via proton-coupled solute transport across the inner mitochondrial membrane. Yet, unlike the cytosolic pH, pH in the mitochondrial matrix did not drop below the pre- illumination levels after the initial alkalinisation peak. Alkalinisation of the mitochondrial matrix after a light period was recently observed, albeit without any pH measurement in the cytosol, and therefore interpreted as increased mETC activity and proton pumping rate leading to a steepened proton gradient across the inner mitochondrial membrane^50,54^. The more comprehensive, high resolution dataset obtained here instead suggests that the matrix pH signature is likely to result as a combination of both, (i) coupling with cytosolic pH (cytosolic pH provides the offset against which the inner membrane proton gradient is established), which dominates initially, and (ii) an increased proton gradient between cytosol/intermembrane space and the mitochondrial matrix, possibly due to photosynthesis-dependent stimulation of mitochondrial electron transport^64^. After illumination, the pH in all three compartments did not recover to pre-illumination baseline within the 10 min of measurement (**Figure 3o**), suggesting that full recovery involves a complex set of mechanisms and requires more time^80^.

### ‘pH wave’ propagation across the cell via proton coupled transmembrane transport

Resolving the physical players and the specific mechanisms that underpin the observed pH connectivity between chloroplasts, cytosol, and mitochondria is not straightforward, since a large set of proton-associated transport processes occur in the membranes and are likely to jointly contribute to the cross-compartment ‘pH wave’. Potential players include, for instance, NHD1 that functions as a potential Na⁺/H⁺ antiporter in the chloroplast inner envelope^94^, Day-length-dependent delayed- greening1 (DLDG1) that acts as a putative K⁺(Ca^2^⁺)/H⁺ antiporter or regulator^95^, Ycf10, a homolog of DLDG1^96^, and fluctuating-Light Acclimation Protein1 (FLAP1) that regulates both DLDG1 and Ycf10 and contributes to proton flux^97^. Further, BIVALENT CATION TRANSPORTER 2 (BICAT2)/ CHLOROPLAST MANGANESE TRANSPORTER1 (CMT1), a putative Mn^2+^ and Ca^2^⁺/H⁺ antiporter, may also contribute^98,99^. While K^+^ exchange antiporters (KEA) 1 and 2 contribute to stromal pH homeostasis^100^, recent studies suggest their involvement in light-induced stroma alkalinization is limited^66^. In mitochondria, the phosphate transporters MPT2 and MPT3, located in the inner mitochondrial membrane, are among the most abundant mitochondrial proteins^101^, facilitating Pi flux in exchange with OH⁻ (equivalent to H⁺ symport) to sustain oxidative phosphorylation^102–104^. Mitochondria may even bypass the cytosol in proton transfer to chloroplasts aided by proximity of their membranes through organelle associations^69^ ^70^.

### Stromal and cytosolic

*ATP levels are elevated in the light, but show distinct signatures.* The dynamics of MgATP^2-^ pools in the chloroplast stroma and the cytosol exhibited synchronization at dark-light transitions, as evidenced by the parallel increase in stromal and cytosolic MgATP^2-^ concentration. The increase in temperature is likely to have a quantitative impact on ATeam1.03-nD/nA measurements due to the temperature sensitivity of the sensor *K*d^78^. Yet, at light fluxes of up to 200 µmol m^-2^ s^-1^ the measured changes in temperature were limited (up to 2°C) (**Supplemental Figure 15c**). Further, the signatures of the sensor responses differed between the two compartments suggesting that the observed responses were not dominated by temperature changes (**Figure 4b, e**). While ATeam1.03-nD/nA is temperature-sensitive, other sensors, such as cpYFP and Peredox, showed minimal sensitivity to temperature changes (**Supplemental Figure 19a, b**). Additionally, the application of DCMU abolished sensor responses, indicating that observed dynamics were directly tied to photosynthesis (**Figure 5f, Supplemental Figure 19c**).

The different MgATP^2-^ signatures in the chloroplast stroma and the cytosol corroborate the current consensus that cytosolic ATP is predominantly of mitochondrial origin, or generated in the cytosol itself^50,64,81^. The initial increase in stroma MgATP^2-^ is driven by photochemical ATP synthesis, which slows as the CBB cycle activates and other ATP consuming processes. Interestingly, and in analogy to pH (**Figure 2c**), higher light intensities than 20 µmol m^-2^ s^-1^ did not change the dynamics of the sensor (**Figure 4b**), suggesting that ATP synthesis is fully operational at low light intensities and increases in steady-state ATP concentrations are constraint, e.g. by buffering through adenylate kinases and proportional activation of consumption. Similar observations were recently made in *Physcomitrium patens*^64^, where the cytosolic ATP increase reached saturation at 50 µmol m^-2^ s^-1^. Following illumination, cytosolic ATP levels continue to increase even after the light is turned off, likely due to light-enhanced dark respiration (LEDR)^105,106^. LEDR, which can sustain elevated respiration rates for up to 20 minutes after illumination^106^, may rely on the remaining reducing power in the cytosol, as indicated by sustained reduction of the cytosolic NAD pool after 10 minutes of illumination (**Figure 5**). Elevated post-illumination ATP levels were previously observed in *Physcomitrium patens*^64^ and Arabidopsis cotyledons^50^. LEDR can be abolished by the loss of mitochondrial complex I activity, highlighting its dependence on mitochondrial respiration^64^. Moreover, the stability of the MgATP^2-^ complex, which is the compound that the ATeam1.03-nD/nA sensor binds, is pH sensitive. Thus, the pH changes during dark-light transitions (**Figure 3**) may in principle contribute to the formation and dissociation of the MgATP^2-^ complex independent of changes in concentration of free ATP or adenylate charge^48^. The MgATP^2-^ signatures observed in the stroma and the cytosol (**Figure 4**) show striking similarity to previous ATP measurements in chloroplast extracts obtained by NAF from *Spinacia oleracea* and *Beta vulgaris* leaves^37^. While the biosensor data provide higher time resolution, the NAF data validate the occurrence of bona fide changes in adenylate charge.

### Cytosolic NAD redox signatures indicate distinct phases of photosynthesis-derived reductant export in vivo

We observed a pronounced, photosynthesis-dependent reduction of the cytosolic NAD pool (**Figure 5**) demonstrating that photosynthetic activity has a major impact on the cytosolic NAD redox state. A similar effect was recently found in another study which assessed the impact of dark-light transitions in Arabidopsis cotyledons expressing SoNar, a different NADH/NAD^+^-sensitive biosensor^53,54^. While several of the NAD redox datasets are in qualitative agreement, there are obvious differences at the quantitative level of the specific signatures. An important factor is likely to be the time delay between illumination and imaging that we minimized through the on-stage illumination setup introduced here. Further, different from Peredox-mCherry which is hardly affected by physiological pH changes, SoNar shows a major pH artefact which is particularly relevant in the context of photosynthetic analyses^107^. While correction routines or alternative imaging protocols have been applied, those are themselves prone to introducing errors. More recently a fusion of SoNar with mCherry has been introduced in which the major pH artefact of SoNar can be circumvented^53^.

The complex NAD redox signature that we observe consistently at the onset of illumination indicates different phases of reducing equivalent export from the chloroplast stroma. Initially, a rapid reduction of cytosolic NAD likely reflects the activity of the malate valve and the TP/3-PGA shuttle.

Cytosolic NAD reduction reaches similar levels within the first minute across a broad range of light intensities (**Figure 5b**), limited either by stromal NAD(P)H accumulation or by shuttle capacity^108^. However, our data indicate that the loss of key organellar malate dehydrogenases involved in the malate valve (NADP-MDH, mMDH1, and mMDH2) does not fully disrupt NAD dynamics at dark-light transitions (**Figure 6a, b**). This finding indicates that mechanisms other than the malate valve are able to facilitate redox balance during photosynthesis onset *in vivo*, which aligns with previous studies that revealed minimal phenotypic impact from NADP-MDH loss^86^. While TP/3-PGA shuttle and possibly photorespiration are likely candidates to compensate for the export of reducing power in principle, their components require time after the onset of illumination to become fully operational. Also, activation of NADP-MDH has been found to require several minutes^87^. The rapid export of reducing power from the stroma within the first 15 s as observed here, suggests that our current understanding of the *in vivo* regulation of the known shuttle systems requires expansion, or the existence of alternative redox shuttles.

Cytosolic NAD undergoes transient oxidation after 1 to 4 minutes of illumination between 20 and 600 µmol m^-2^ s^-1^ (**Figure 5b, c**). Activation of the CBB cycle and ATP synthase, as well as of photorespiration after the first minutes of illumination, are plausible mechanisms. The oxidation is likely to be a result of reduced export rates from the chloroplast (even re-import of reducing power appears plausible) as the CBB cycle increases its capacity as a sink for stromal NADPH, and the transfer of reducing equivalents to the peroxisome as photorespiration ramps up^16^. After the short oxidation, the cytosolic NAD pool is further reduced in a light intensity-dependent manner (**Figure 5b, d**), probably reflecting stable CBB cycle activation, while an excess of reducing equivalents appears to be generated by photosynthetic electron transport which is exported from the stroma. The NAD redox signatures revealed through illumination imaging resolve phases, such as the transient oxidation, that had hitherto been impossible to detect. Yet, interestingly, NAF-based analysis of *Beta vulgaris* leaf cells showed changes in NADH levels in both the chloroplast fraction, as well as the nonchloroplastic part of the cell, that followed a very similar pattern^38^.

### Impairing organelle malate metabolism leads to re-allocation of reducing equivalents between cell compartments

A key role that has been proposed for the mMDH1 and mMDH2 in the light is to keep the NAD pool of the mitochondrial matrix sufficiently oxidized to maintain photorespiratory glycine oxidation^10985^. Interestingly, loss of function of *mmdh1*, the dominant isoform of mMDH, leads to a loss of NAD reduction during illumination (consistently at both 20 and 200 µmol m^-2^ s^-1^), indicating a constraint for less reducing power export from mitochondria (**Figure 6g, h**). The differences in the amplitude of NAD reduction of the three organellar *mdh* lines were found to be mainly due to a more reduced cytosolic NAD pool in the dark (**Figure 6c, d**). Such a major change in cytosolic NAD redox status pinpoints a pronounced re-arrangement of the subcellular metabolic flux landscape. Indeed, constrained mMDH capacity is expected to not just impact malate valve activity, but also other metabolic fluxes. When the TCA cycle runs in its cyclic mode in the dark limiting mMDH capacity is likely to force malate export from the mitochondrial matrix to the cytosol, which may explain the observed shift of the cytosolic NAD redox state to a more reduced state (**Figure 6e, f**). Similarly, constraint mMDH capacity may inhibit the import of malate released from the vacuole, further increasing malate levels in the cytosol, which can be oxidized to OAA by cytosolic MDH, thereby reducing the cytosolic NAD pool^110^. Recent findings in tobacco^111^ support the notion that a major re-allocation of reductant between cell compartments by re-routing malate fluxes may be a central feature of the metabolic redox network of photosynthetic plant cells. Interestingly, reduction of the cytosolic NAD pool was also found in the *nadp-mdh* line. The observation of such a pronounced impact of the lack of NADP-MDH even in the dark was surprising considering that NADP-MDH activation requires the light-dependent reduction via the stromal thioredoxin system^87^. While it is difficult to rule out the possibility that stromal NADP-MDH may have residual activity in the dark *in vivo*, the impact on cytosolic NAD redox status may be less direct. For instance, loss of NADP-MDH has been found to induce the expression of plastid-located NAD-dependent malate dehydrogenase, and glyceraldehyde-3-phosphate dehydrogenase in the dark^112^. Hence, the interpretation of re-wired reductant partitioning deserves a ‘systems’ perspective that may be achieved through a combination with state-of-the-art omics approaches in the future. This and similar questions may now be systematically addressed by applying and adjusting the advanced illumination imaging technology introduced here. We anticipate the combination of live environmental control, a growing repertoire of biosensors, the spectroscopic analyses of the photosynthetic pigments and the use of reverse genetic approaches^88,89,113,114^, to be a powerful driver of discovery and dissection of the mechanisms that underpin photosynthetic physiology and acclimation across the entire plant cell, across tissues and even organs like whole leaves.

## Materials and Methods

### Plant material and growth conditions

*Arabidopsis thaliana* (L.) Heynh. (accession Columbia-0, Col-0) seedlings were grown on half-strength (0.5x) Murashige and Skoog medium^115^ under long-day conditions (16 h light, 75-100 µmol s^-1^ m^-2^, 22 °C and 8 hours dark, 18 °C) after stratification at 4 °C for two days. To obtain discs of mature rosette leaves, Arabidopsis plants were grown for 4 to 5 weeks in imbibed Jiffy-7 pellets (Jiffy Products International BV, Zwijndrecht, Netherlands) under long-day conditions (100-120 µmol s^-1^ m^-2^). The fluorescent protein biosensors cpYFP, pH-GFP, ATeam1.03-nD/nA and Peredox-mCherry were stably expressed in different subcellular compartments in *Arabidopsis thaliana* as described before^44,46,48,52,56^. All experiments were carried out on true leaves from 4- to 5-week-old Arabidopsis plants unless specified.

### Generation of plant lines

A plasmid encoding Peredox-mCherry under the control of the *Ubiquitin-10* promotor^52^ was used to stably transform the homozygous insertional lines *mmdh1* (GABI_097C10) and *mmdh2* (SALK_126994)^85^, and *nadp-mdh* (SALK_012655)^86^ using the floral dip method^116^. Positive transformants were identified by selection on hygromycin B and at least three independent lines per genotype were isolated based on their fluorescence intensity. Two (*nadp-mdh*, *mmdh2*) or three (*mmdh1*) independent sensor lines were used for time series acquisition. Transformants displayed no relevant phenotype compared to their non-transformed mutant background counterparts.

### Automated illumination system

Dynamics at dark-to-light transitions were monitored using a Zeiss LSM 980 microscope equipped with a 10x lens (Plan-Apochromat, 0.3 N.A.) (Carl Zeiss Microscopy GmbH, Jena, Germany). cpYFP localisation was performed using a Zeiss LSM 980 using the 40x lens (C-Apochromat, 1.2 N.A.). The customised on-stage illumination system was connected to the LSM 980 via the Zeiss trigger interface, enabling to control of the microscope and the illumination system in a coordinated manner using the Experiment Designer in ZEN blue 3.5. A custom-built device (Electronics workshop, Faculty of Chemistry, University Bonn, Germany) uses the 5 V trigger signal to switch the cold-white LED (Max Pferdekaemper GmbH & Co. KG, Menden, Germany) implemented in the on-stage illumination system (**Supplement Figure 2**).

### Confocal laser scanning microscopy

Confocal imaging were carried out on fully expanded leaves from 4 to 5-week-old Arabidopsis plants before flowering as described previously^117^ unless specified. To minimise the barrier between the light source and sample, each leaf disc was mounted between two coverslips (18 x 18 mm and 22 x 40 mm, VWR International GmbH, Darmstadt, Germany). cpYFP and pH-GFP fluorescence were excited at 405 nm and 488 nm, while emission was collected at 508-535 nm. To control for autofluorescence, emission after excitation with 405 nm was simultaneously recorded at 430-470 nm. ATeam1.03-nD/nA was excited at 445 nm and emission was recorded at 466-499 nm (CFP) and 526-561 nm (Venus). Peredox-mCherry fluorescence was excited at 405 nm (tSapphire) and 561 nm (mCherry) and emission was recorded at 499-544 nm and 588-624 nm, respectively. Autofluorescence was collected at 430-451 nm. For measurements with all three sensors, chlorophyll fluorescence was collected with excitation at 488 nm and emission at 651-700 nm. The pinhole was set to 96 µm (cpYFP), 141 µm (ATeam1.03-nD/nA) and 145 µm (Peredox-mCherry). All plants, which were used for measurements during the same day, were transferred from the growth chamber at once and dark-adapted for at least one hour prior to time series acquisition. Further microscopic settings and details on the illumination regime are given in **Supplemental Table 1**.

### Ratiometric analysis and statistics

The confocal microscopy time series datasets were processed with a custom MATLAB-based software^118^ using x,y noise filtering. The pH, MgATP^2-^ and NADH/NAD^+^ levels are indicated by the ratio of separate channels of sensors. Ratios were then log10-transformed before statistical analysis was performed using GraphPad Prism (version 8.0.1, GraphPad Software, San Diego, CA, USA). In cases where the data were normalized, the mean of the last 5 data points acquired before illumination was used as the reference value. For dark control samples, the data points in the corresponding time window were used for normalization.

### Calibration of cpYFP

A protocol is adopted from Wagner, 2019. 100 mg leaf material was collected and disrupted by TissueLyser II (Qiagen, Hilden, Germany) at 30 Hz for 2 min with pre-cool racks. A cell extract was prepared by adding 350 µl calibration buffers with different pH. After 10 to 30 s vortex, stand the mixture on ice for 5 min. Calibration buffers contained 100 mM NaCl, 1 mM Na2EDTA and for pH values between 5.5 and 6.5 either 100 mM 2-(N-morpholino) ethanesulfonic acid (MES), or between 7.0 and 8.2 100 mM HEPES. Apply the same method to prepare slides for the illumination system, only replace leaf discs with cell extract on the upper layer. Run 3 min time series experiment with cpYFP acquisition setting in the dark, and pick the last data point to represent the sensor ratio in that pH.

### Chlorophyll fluorescence analysis

A Maxi Imaging PAM chlorophyll fluorometer (Heinz Walz, Effeltrich, Germany) was used for chlorophyll fluorescence analysis as previously described^119^ and NPQ was calculated as detailed recently^120^.

## Acknowledgements

We thank Ilka Haferkamp (Kaiserslautern), Beatriz Moreno García (Oxford), Karl-Josef Dietz (Bielefeld) and Philippe Fuchs (Münster) for fruitful discussions, Jörg Imbrock (Münster) and Simon Laubrock (Münster) for help with the spectral measurements of the LEDs, Mark Fricker (Oxford) for lomg-term support with custom image analyses, and Luis Garcia Rodriguez (Münster) support in executing the engineering and optimization of the illumination system. We further are grateful to Thomas Zobel and Sarah Weischer and the Imaging Network of the University of Münster (RI_00497).

## Author contributions

M.S. designed the research; K.Z., M.E., E.F.A, F.K. and F.B. performed the experiments; K.Z., M.E., E.F.A, J.O.N., P.B., A.C., M.H., J.G., M.P.M. and F.B. analysed the data; K.Z., M.E., S.W. and M.S. contributed novel analytical tools; M.S. supervised the research with specific input by S.W., J.M.U., M.H., A.J.M., H.H.K., U.A., V.G.M., I.F., M.S.R.; K.Z., M.E. and M.S. wrote the manuscript with input from all co-authors.

## Supplemental Figure Legends

**Supplemental Figure 1.**
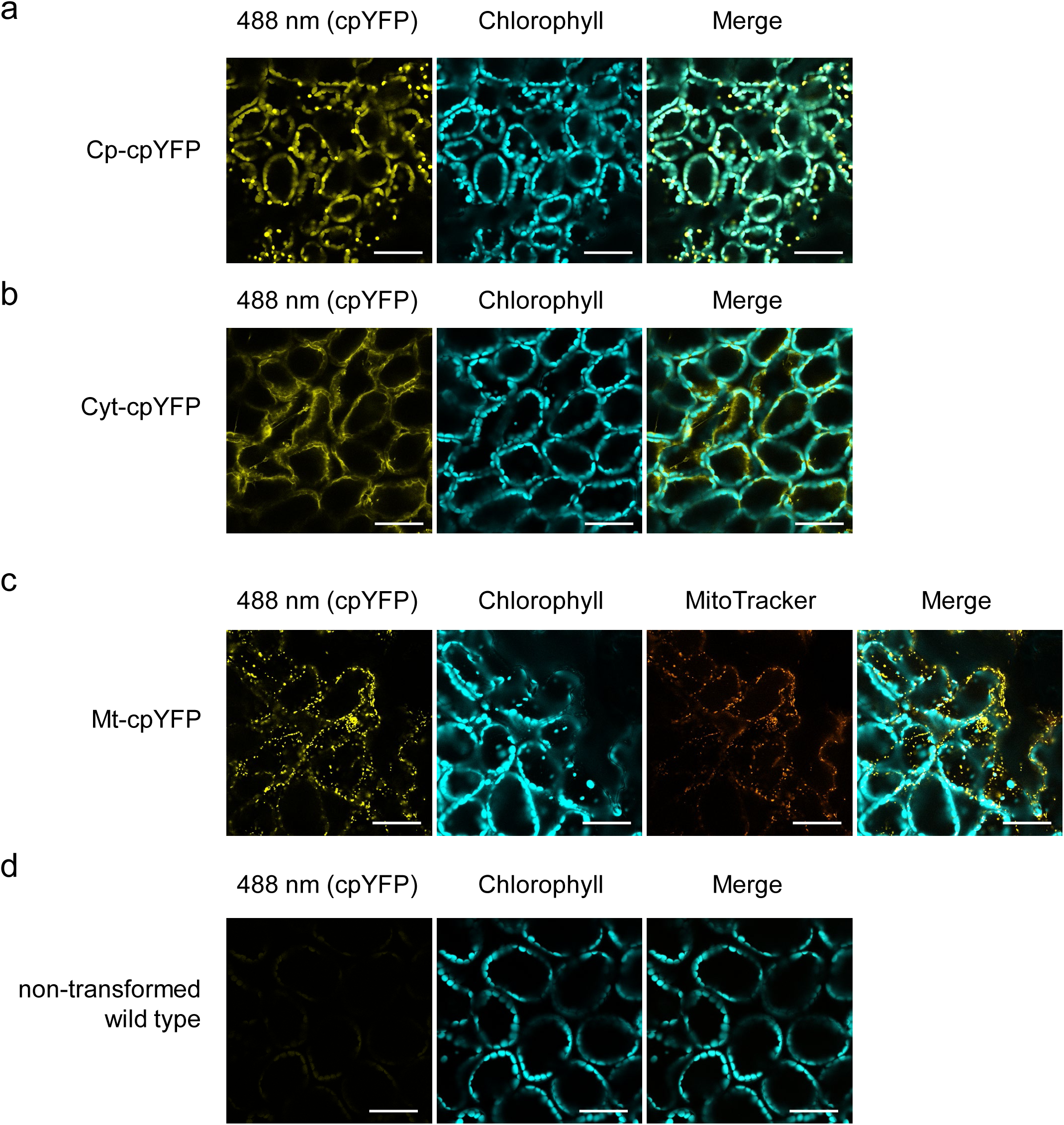
C**o**nfocal **microscopy images of cpYFP localization in the chloroplast stroma, cytosol and mitochondria.** Confocal microscopy images of subcellular localization of cpYFP in the chloroplast stroma **(a)**, cytosol **(b)** and mitochondria **(c)** in mesophyll cells of 4 to 5-week-old Arabidopsis plants. Leaf discs were incubated in 1:2000 dilution MitoTracker^TM^ Orange CMTMRos (MitoTracker) for 20min in **(c)**. **d** Non-transformed wild type imaged with the same confocal settings as in **(a)-(c)**. Confocal setting: excitation: 488 nm; emission cpYFP: 508-535 nm; emission chlorophyll: 651-700 nm; MitoTracker: excitation: 561 nm, emission: 570-579 nm. Scale bars: 50 µm.

**Supplemental Figure 2.**
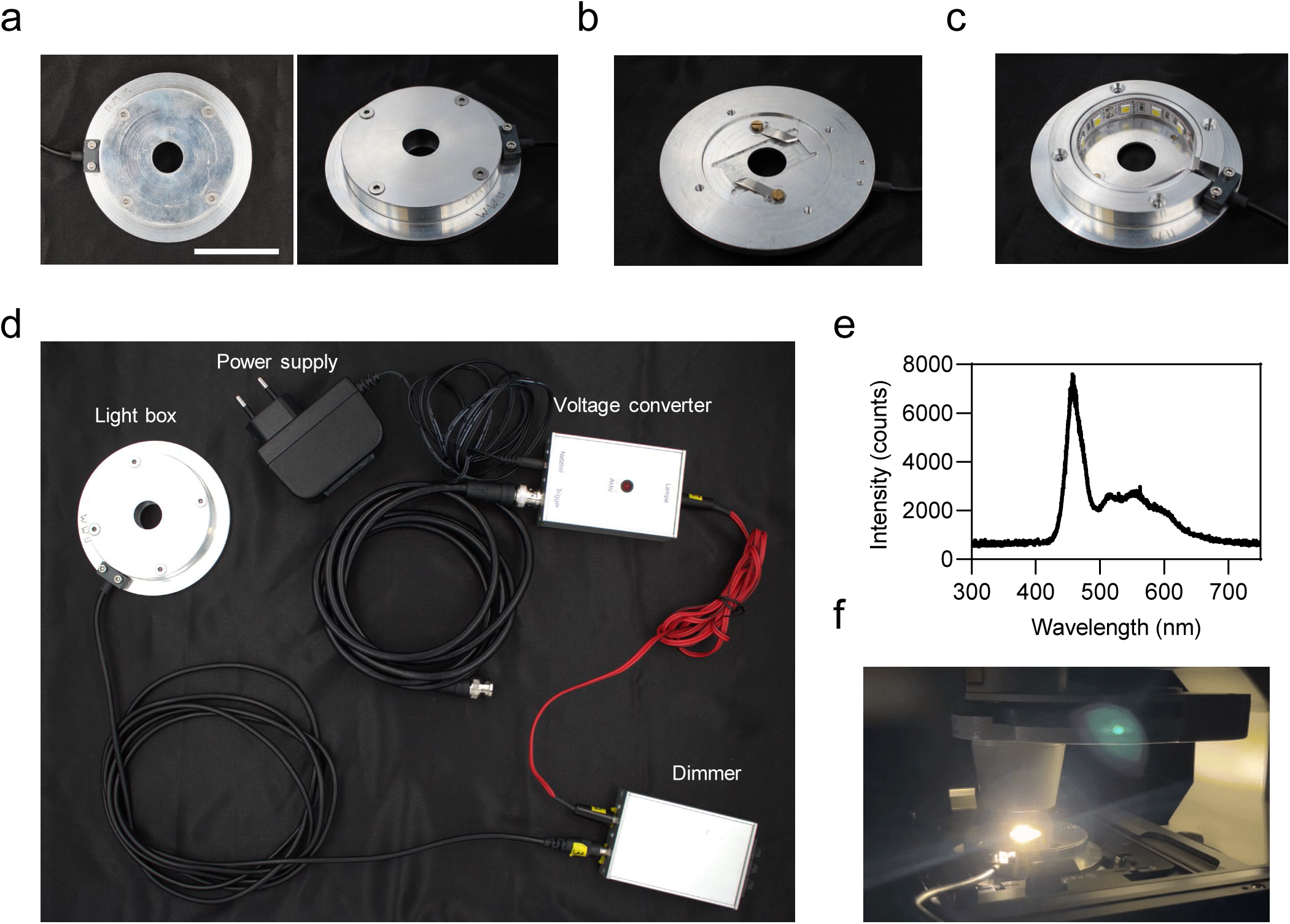
Automated illumination system for confocal laser scanning microscopy. The top **(a)** and bottom **(b)** view of the light chamber. **c** A cold white LED stripe as light source. **d** Assembled light chamber and electrical components to control light intensities. **e** Spectrum of the implemented LED strip recorded with a high-resolution spectrometer (Ocean Optics Inc., Largo, FL, USA). **f** Illumination of a sample during time series acquisition. Scale bar: 5 cm.

**Supplemental Figure 3.**
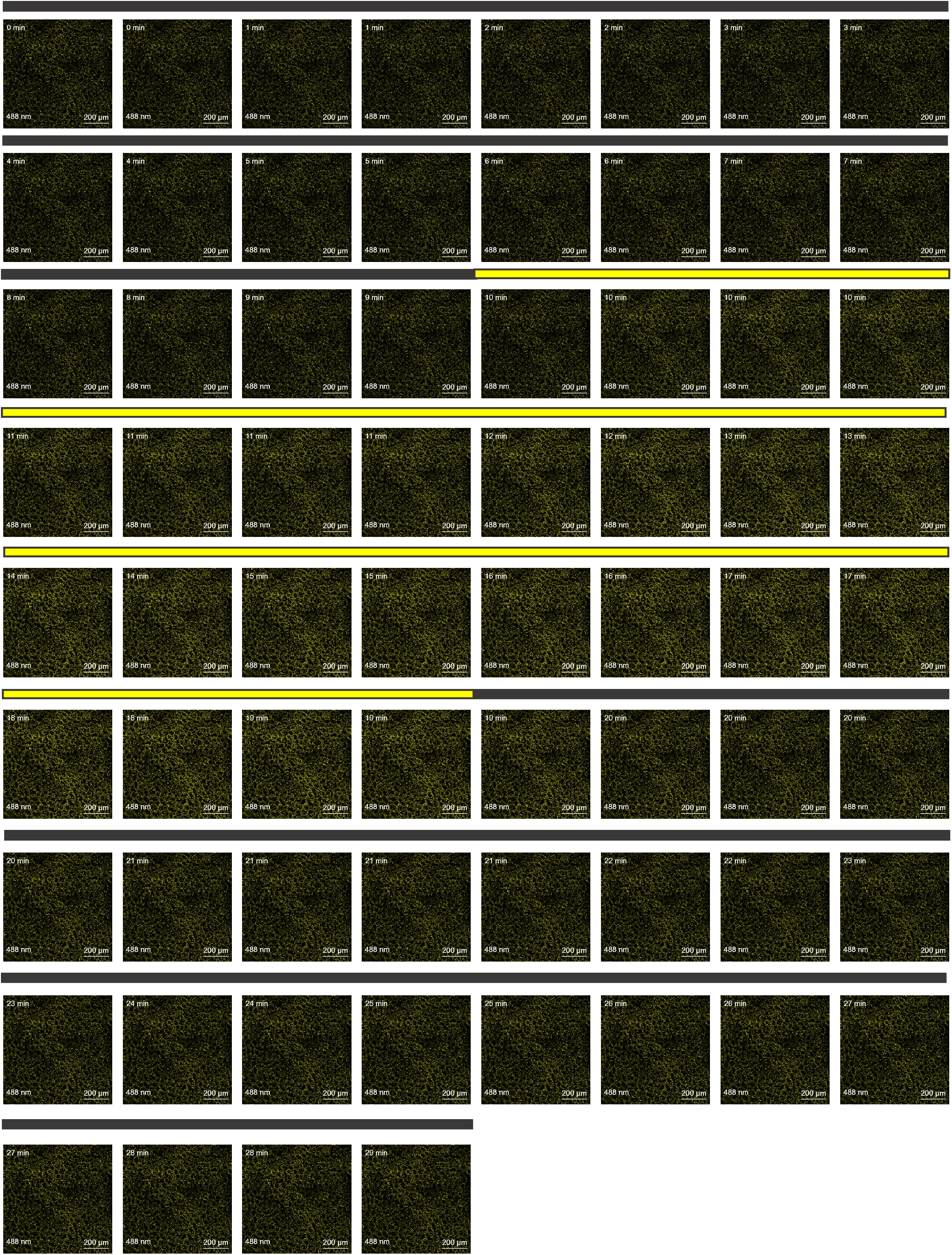
Confocal microscopy images of chloroplast stroma- located cpYFP in dark-light transitions. Confocal images of chloroplast stroma-located cpYFP at 488 nm excitation and 508- 535 nm emission under dark-light transitions (60 µmol m^-2^ s^-1^). The emission intensity (shown in yellow) at 488 nm excitation indicates the extent of alkalinisation in the chloroplast stroma. The black and yellow bars show the dark and light phases separately.

**Supplemental Figure 4.**
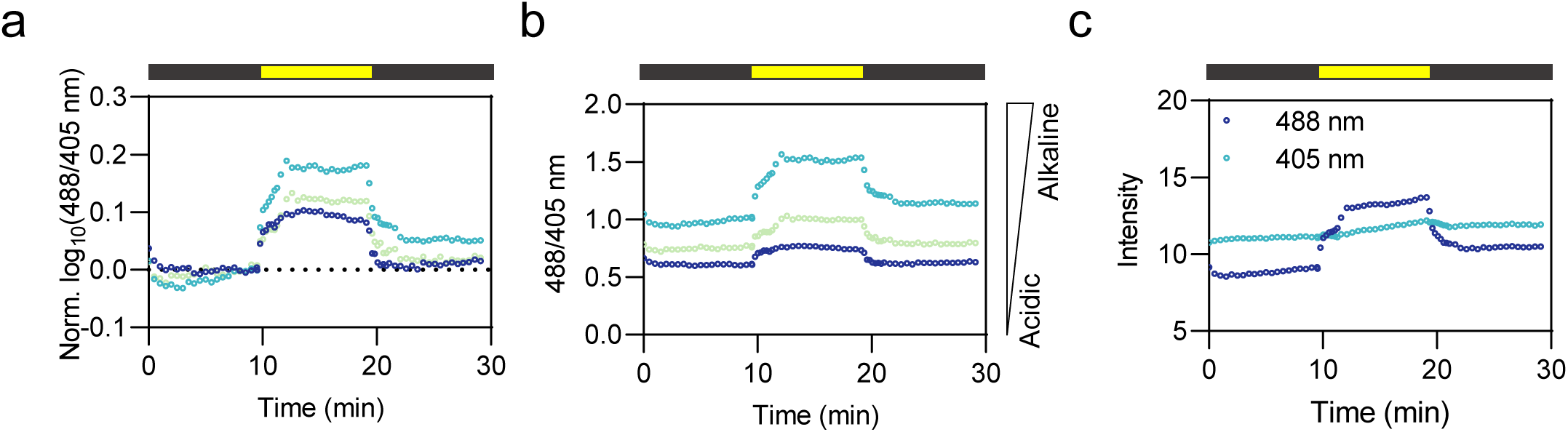
Single replicates and individual channels of chloroplast stroma-located cpYFP under dark-light transitions (60 µmol m^-2^ s^-1^). **a, b** Three replicates of chloroplast stroma-located cpYFP showed similar dynamics under dark-light transitions. In **(a)**, data are normalized to the average of the last five data points prior to illumination and log10-transformed. In **(b)**, data are not normalized or log10-transformed. **c** Emission of single channels of stroma-located cpYFP in response to dark-light transitions. The black and yellow bars show the dark and light phases separately.

**Supplemental Figure 5.**
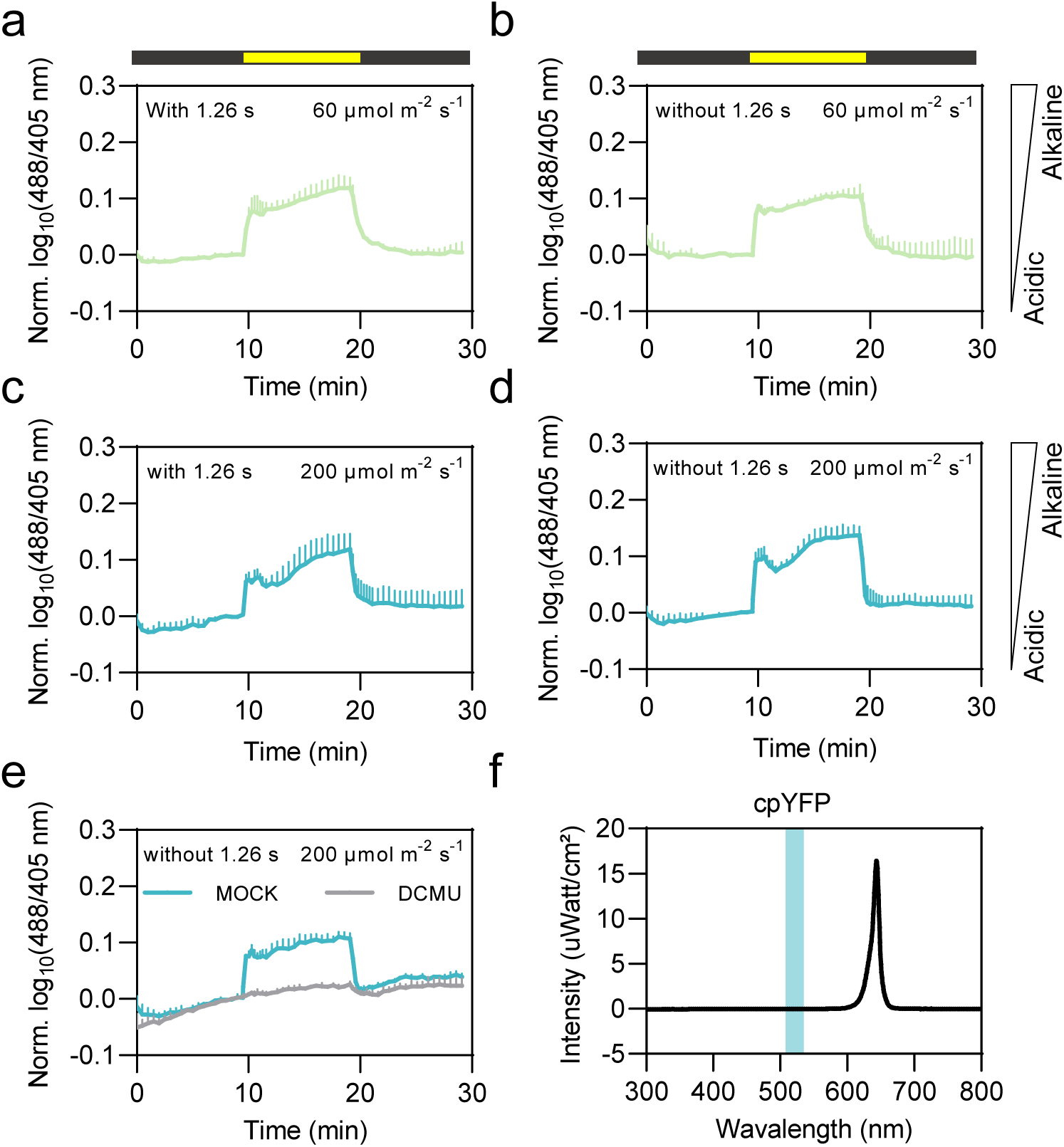
Chloroplast stroma pH dynamics assessed by illumination imaging using red LED illumination The effect of 1.26 seconds interval to pH dynamics was tested under the illumination of the red LED. pH in the chloroplast in response to 60 (green) and 200 (cyan) µmol m^-2^ s^-1^ red LED light (indicated by yellow bars) for 10 min with 1.26 seconds intervals is shown separately in **(a)** and **(c)**, without 1.26 seconds intervals (continuous light) is shown in **(b)** and **(d)**. In **(e)**, ’DCMU’ control of chloroplast stroma located cpYFP: Leaf discs were pre-incubated in 20 µM DCMU and 0.2% (v/v) EtOH, or 0.2% (v/v) EtOH, respectively, for 35 to 45 min. n = 3. Light intensity was 200 µmol m^-2^ s^-1^. Data are normalized to the average of the last five data points in the first dark phase, log10- transformed and averaged. Error bars = SD. **f** Spectrum of the implemented red LED strip recorded with a high-resolution spectrometer (Ocean Optics Inc., Largo, FL, USA). The cyan patch represents the collected emission of cpYFP in the confocal microscope.

**Supplemental Figure 6.**
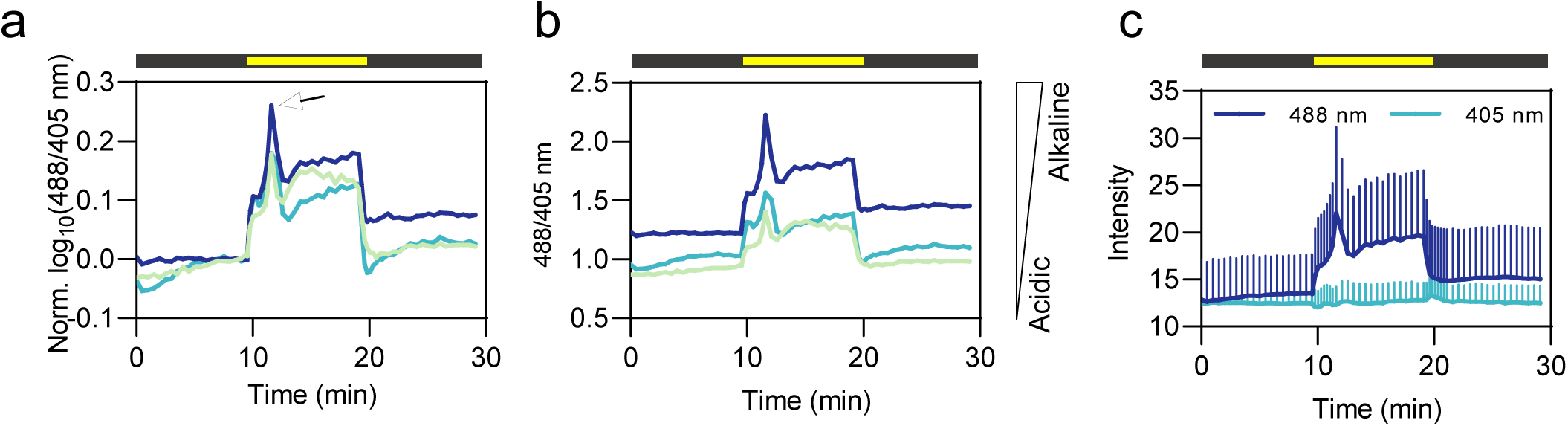
Single replicates and individual channels of chloroplast stroma-located cpYFP ratio at dark-high light transitions (600 µmol m^-2^ s^-1^). Alkalinisation spike (black arrow) shortly after the transition from dark to high light (600 µmol m^-2^ s^-1^) occurred in all individual replicates. In **(a)**, data are normalized to the average of the last five data points prior illumination and log10-transformed. In **(b)**, data are not normalized or log10-transformed. **c** Emission of single channels of stroma- located cpYFP in response to dark-light transitions. The black and yellow bars show the dark and light phases separately. Error bars = SD.

**Supplemental Figure 7.**
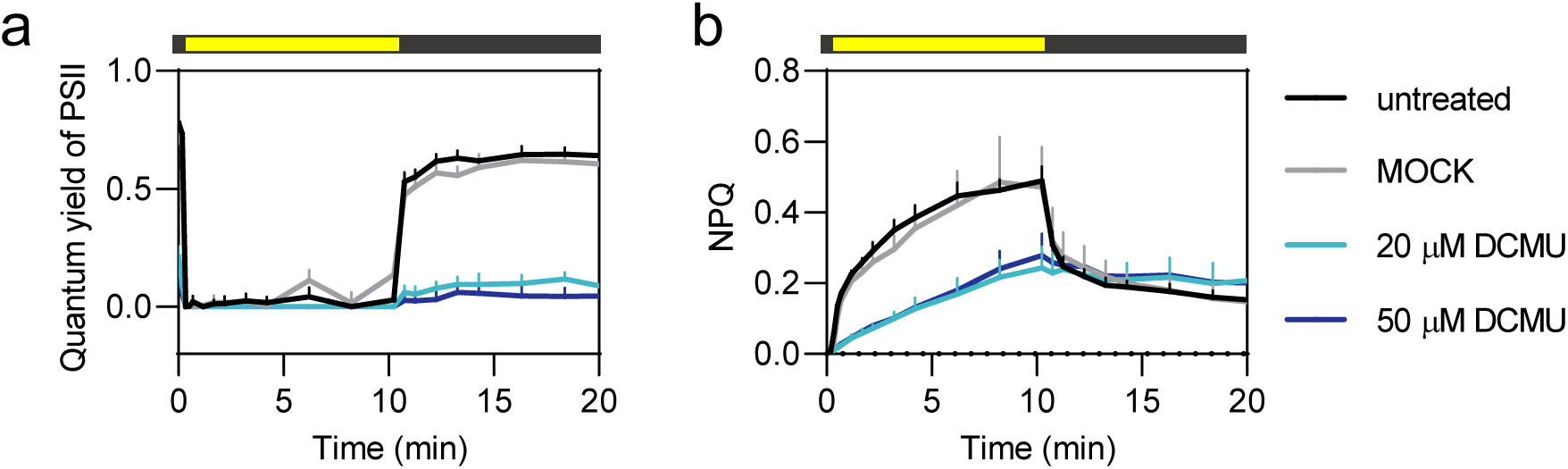
Assessment of the effectivity of DCMU treatments by PAM measurements. The dynamics of **(a)** quantum yield of photosystem II (PSII) and **(b)** non-photochemical quenching (NPQ) were determined in discs of mature leaves of 4 to 5-week-old Arabidopsis plants without biosensor expression. Yellow bars indicate illumination period (65 µmol m^-2^ s^-1^ emitted by the 450 nm LED source of a Maxi Imaging PAM chlorophyll fluorometer; Walz, Germany); black bars indicate the absence of illumination. For treatments leaf discs were pre-incubated in the solvent control (‘MOCK’), 20 µM DCMU and 50 µM DCMU, respectively, for 35 to 45 min in the dark before the measurements were started. n = 8. Error bars = SD. NPQ: non-photochemical quenching.

**Supplemental Figure 8.**
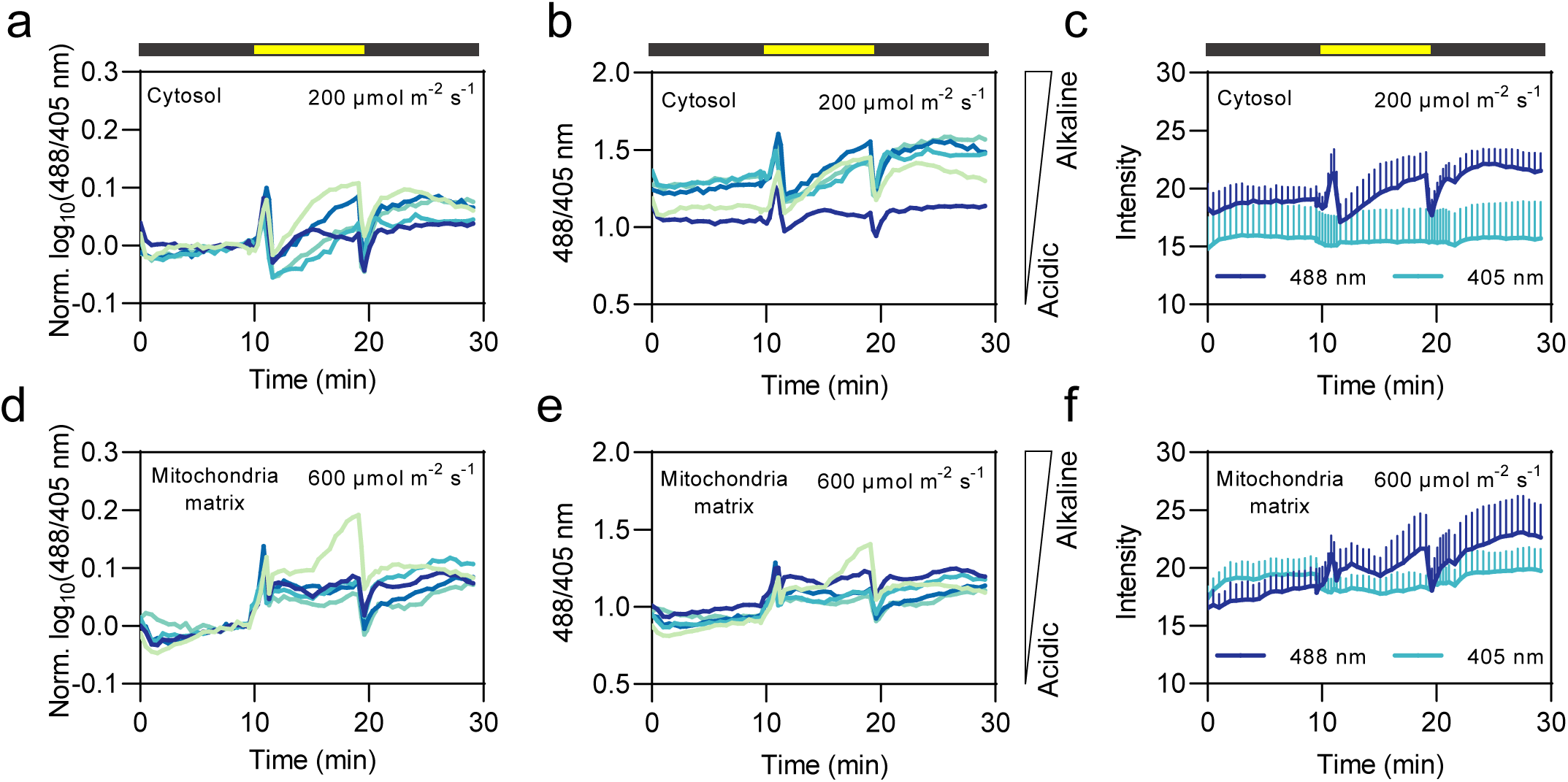
Single replicates and two separate channels of cytosol- and mitochondria-located cpYFP ratio under dark-light transitions. In **(a)** and **(d)**, data are normalized to the average of the last five data points prior to illumination and log10-transformed. In **(b)** and **(e)**, data are not normalized or log10- transformed. Emission of single channels of cytosol- **(c)** and mitochondria- **(f)** located cpYFP in response to dark-light transitions. The black and yellow bars show the dark and light phases separately. Error bars = SD.

**Supplemental Figure 9.**
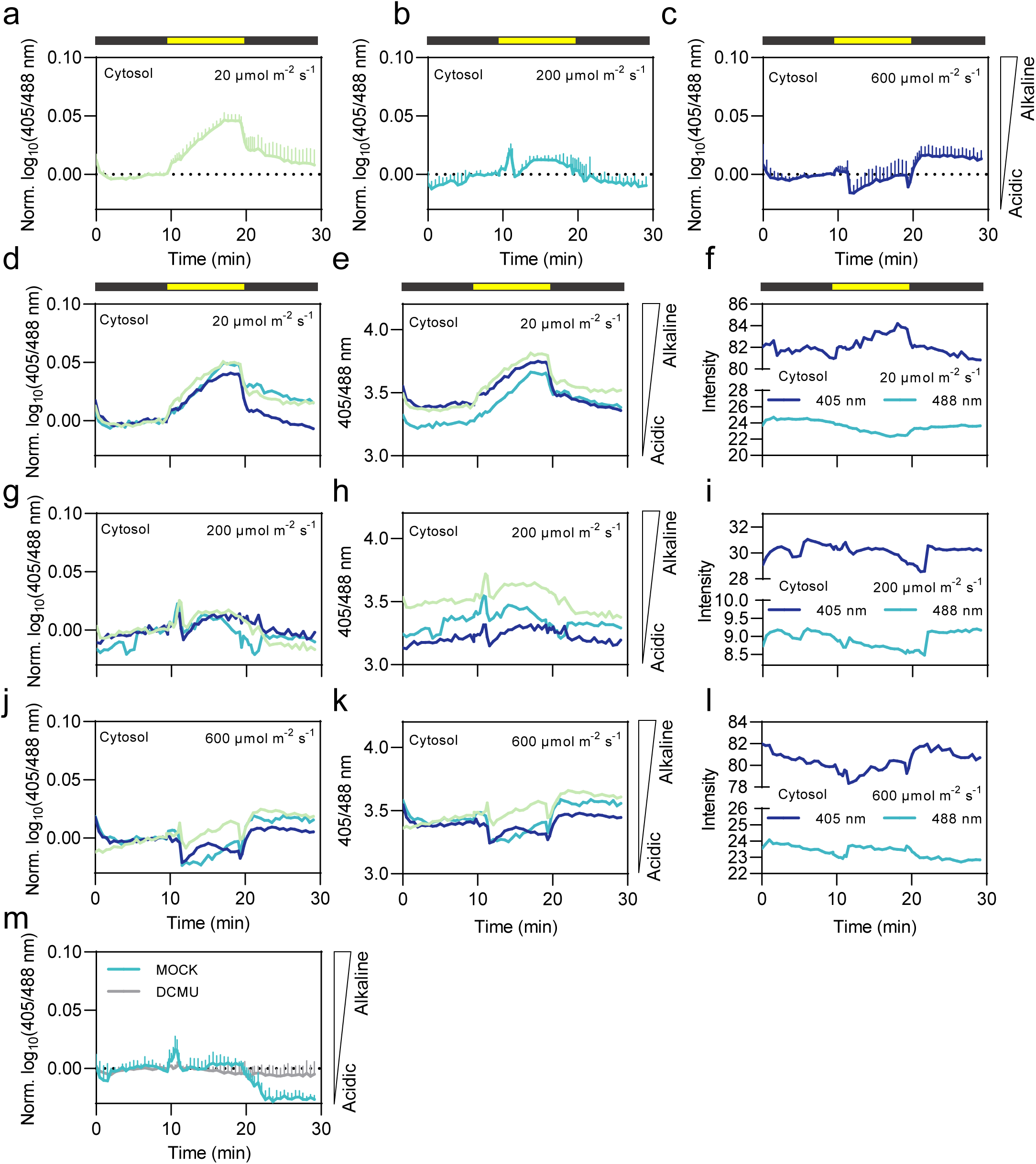
Cytosolic pH dynamic under illumination revealed by pH-GFP Cytosolic pH dynamics in response to 20 **(a)**, 200 **(b)** and 600 **(c)** µmol m^-2^ s^-1^ light (indicated by yellow bars) for 10 min was monitored in discs of mature leaves of 4 to 5-week-old Arabidopsis plants expressing the pH-sensitive fluorescent protein pH-GFP. Single replicates and channels are shown in **(d-l).** In **(d), (g)** and **(j)**, data are normalized to the average of the last five data points prior illumination and log10- transformed. In **(e), (h)** and **(k)**, data are not normalized or log10-transformed. Emission of single channels of pH-GFP in response to dark-light transitions are shown in **(f), (i)** and **(l)**. **m** ‘DCMU’ control and the respective ‘MOCK’ treatment in the cytosol-located pH-GFP. Leaf discs were pre-incubated in 20 µM DCMU and 0.2% (v/v) EtOH, or 0.2% (v/v) EtOH, respectively, for 35 to 45 min. Light intensity was 200 µmol m^-2^ s^-1^. Data are normalized to the average of last five data points prior illumination, log10-transformed and averaged. Error bars = SD. The dotted horizontal lines indicate log10(405/488 nm) = 0. n = 3.

**Supplemental Figure 10.**
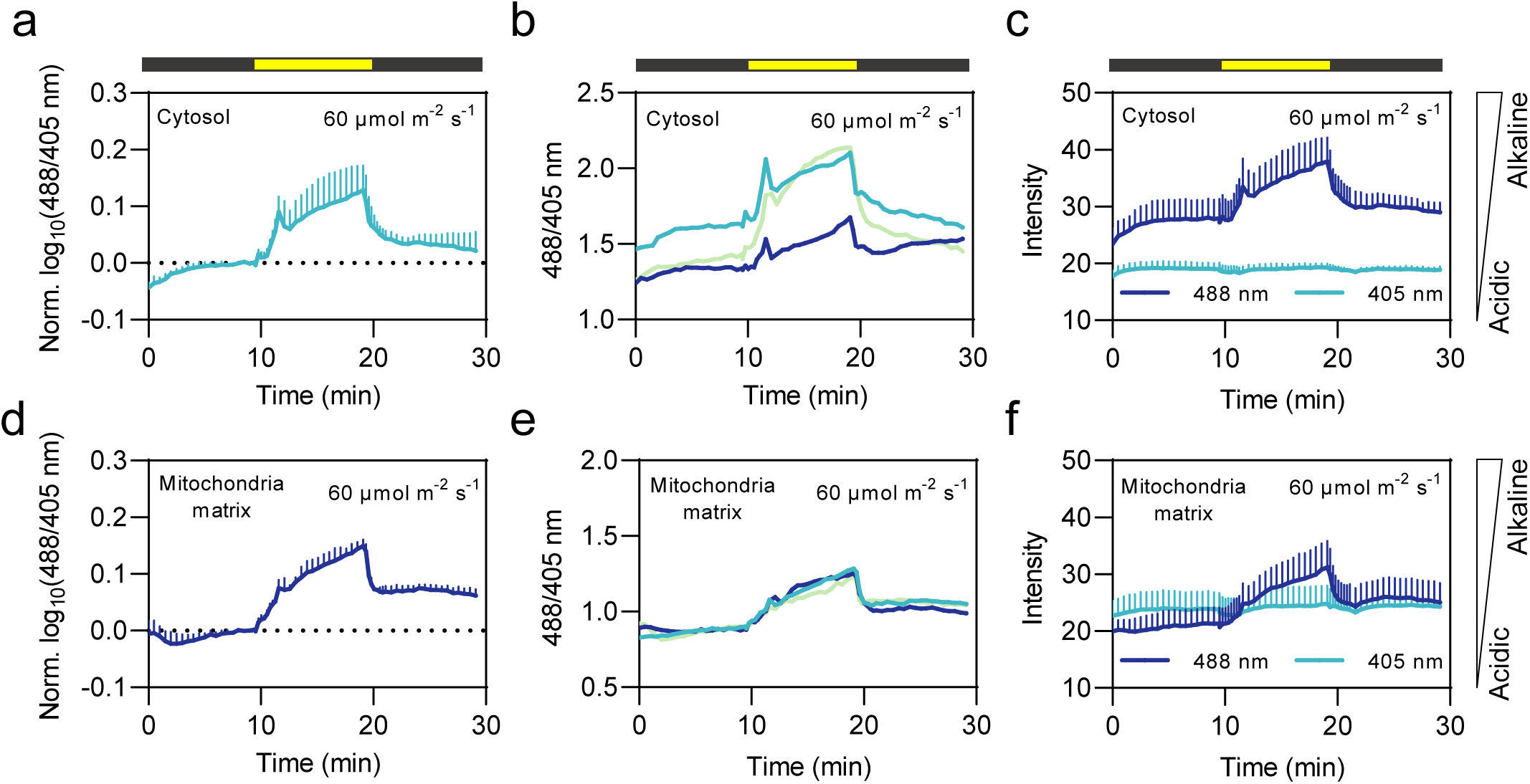
Single replicates and individual channels of cytosol- and mitochondria-located cpYFP ratio under dark-light transitions (60 µmol m^-2^ s^-1^). In **(a)** and **(d)**, data are normalized to the average of the last five data points prior to illumination and log10-transformed. n = 3. In **(b)** and **(e)**, data are not normalized or log10-transformed. Different colors indicate single replicates. Emission of single channels of cytosol- **(c)** and mitochondria- **(f)** located cpYFP in response to dark-light transitions. The black and yellow bars show the dark and light phases separately. Error bars = SD.

**Supplemental Figure 11.**
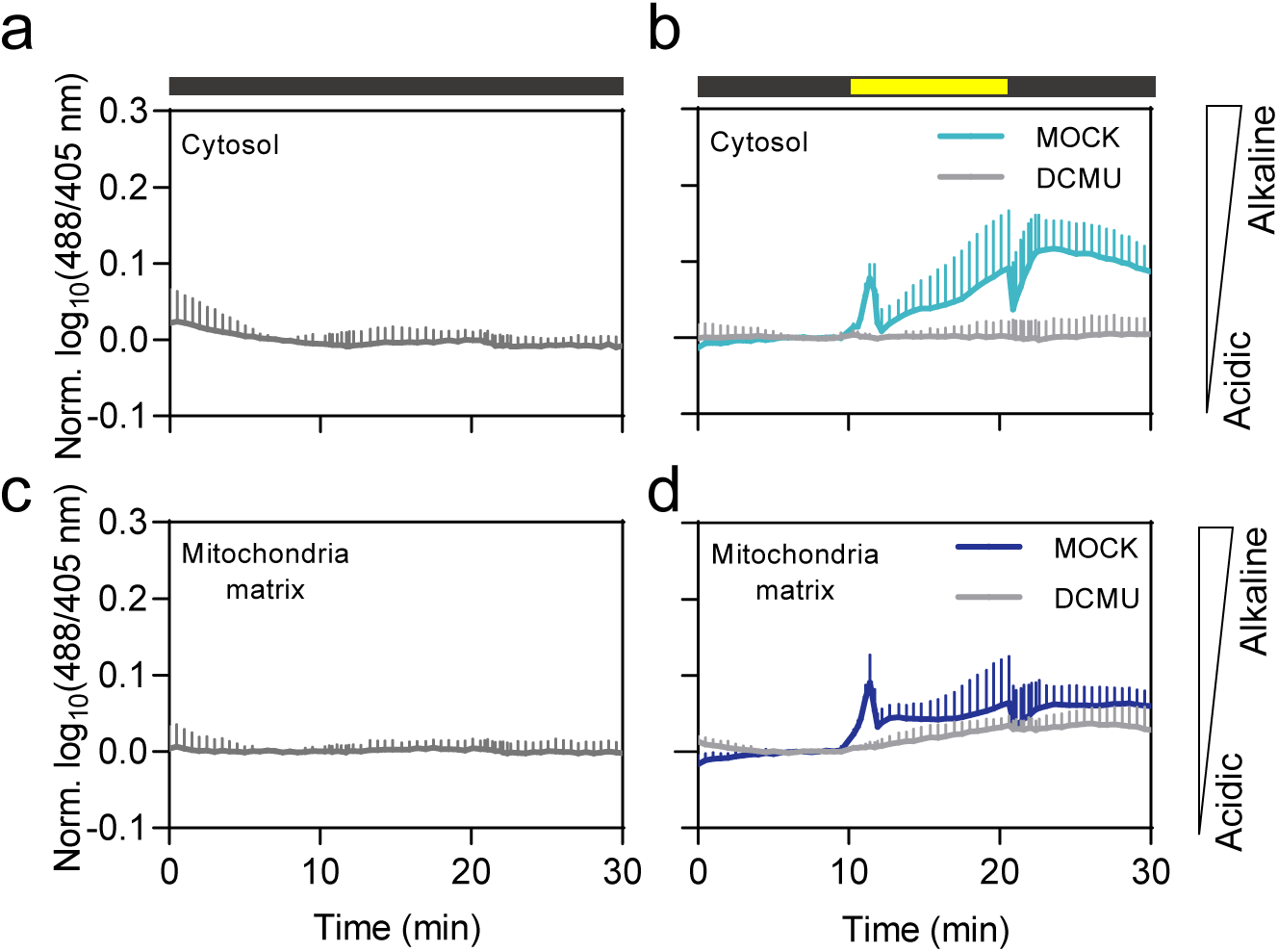
‘No light’ and ’DCMU’ control of cytosol and mitochondria-located cpYFP. ‘No light’ control to cytosol- **(a)** and mitochondria- **(c)** located cpYFP. n = 4. ’DCMU’ control of cytosol- **(b)** and mitochondria- **(d)** located cpYFP: Leaf discs were pre-incubated in 20 µM DCMU and 0.2% (v/v) EtOH, or 0.2% (v/v) EtOH, respectively, for 35 to 45 min. n = 3-4. Light intensity is 200 µmol m^-2^ s^-1^. Data are normalized to the average of last five data points in the first dark phase, log10-transformed and averaged. Error bars = SD.

**Supplemental Figure 12.**
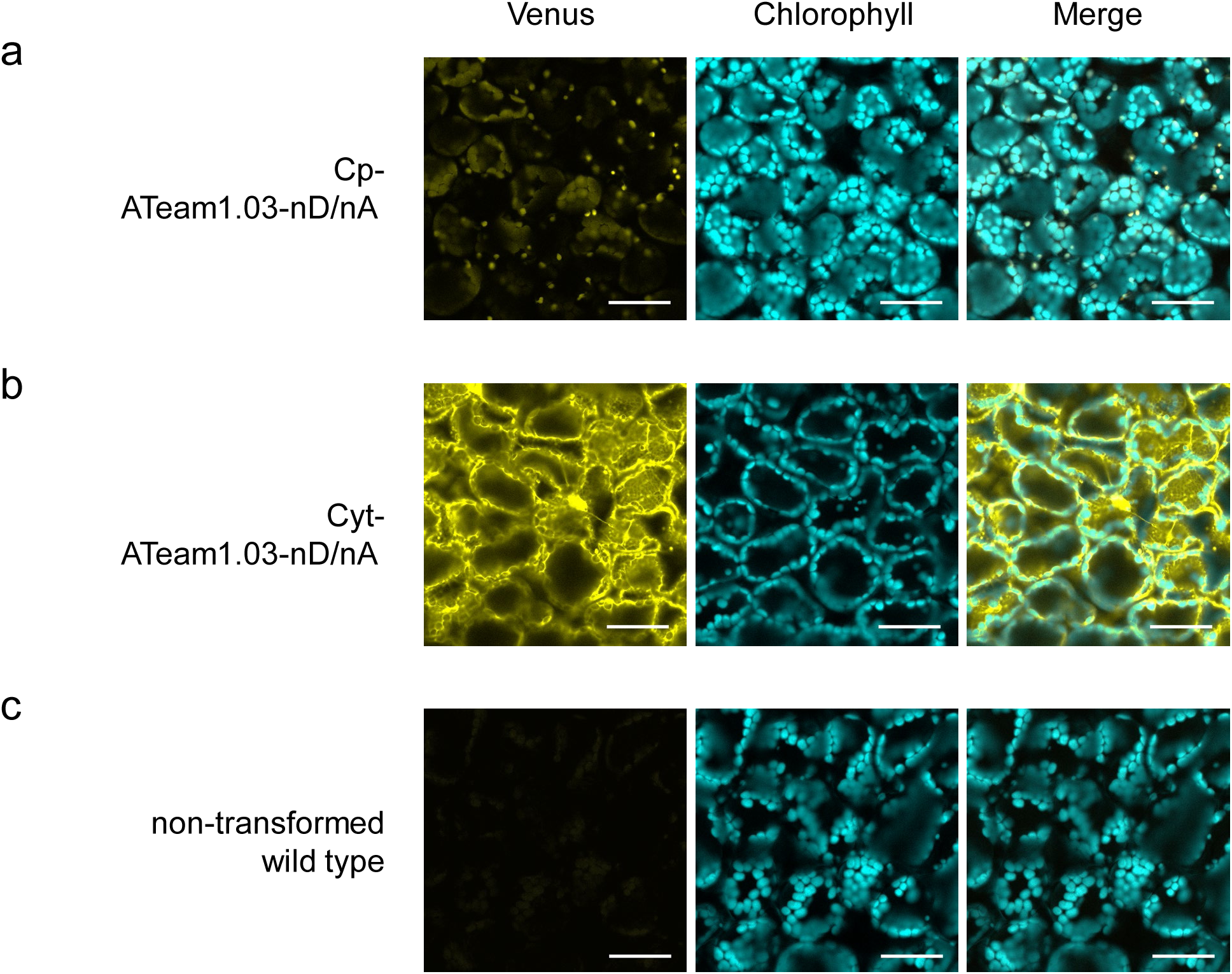
Confocal microscopy images of subcellular localization of ATeam1.03-nD/nA in the chloroplast stroma and cytosol. Confocal microscopy images of subcellular localization of ATeam1.03-nD/nA in the chloroplast stroma **(a)** and cytosol **(b)** in mesophyll cells of 4 to 5-week-old Arabidopsis plants. **c** Non-transformed wild type imaged with the same confocal settings as in **(a)** and **(b)**. Confocal setting: excitation: 445 nm; emission Venus: 526-561 nm; emission chlorophyll: 648-676 nm. Scale bars: 50 µm.

**Supplemental Figure 13.**
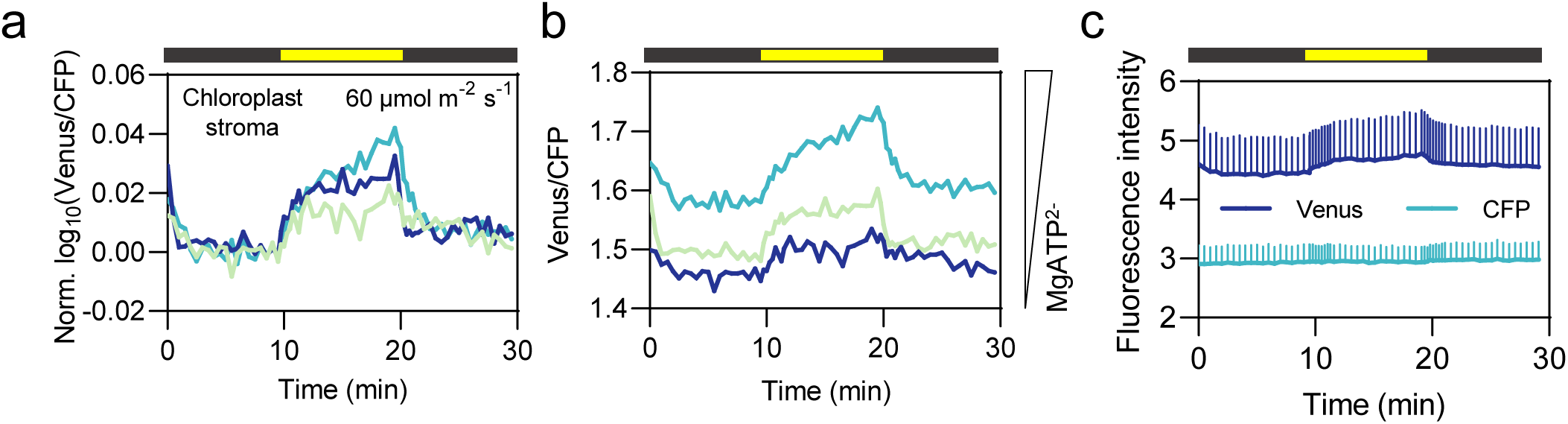
Single replicates and individual channels of chloroplast stroma-located ATeam1.03-nD/nA under dark-light transitions (60 µmol m^-2^ s^-1^). **a, b** Three replicates of chloroplast stroma-located ATeam1.03-nD/nA showed increasing MgATP^2-^ under 60 µmol m^-2^ s^-1^. In **(a)**, data are normalized to the average of the last five data points prior illumination and log10-transformed. In **(b)**, data are not normalized or log10-transformed. Different colors indicate single replicates. **c** Emission of single channels of stroma-located ATeam1.03-nD/nA in response to dark-light transitions. The black and yellow bars show the dark and light phases separately. Error bars = SD.

**Supplemental Figure 14.**
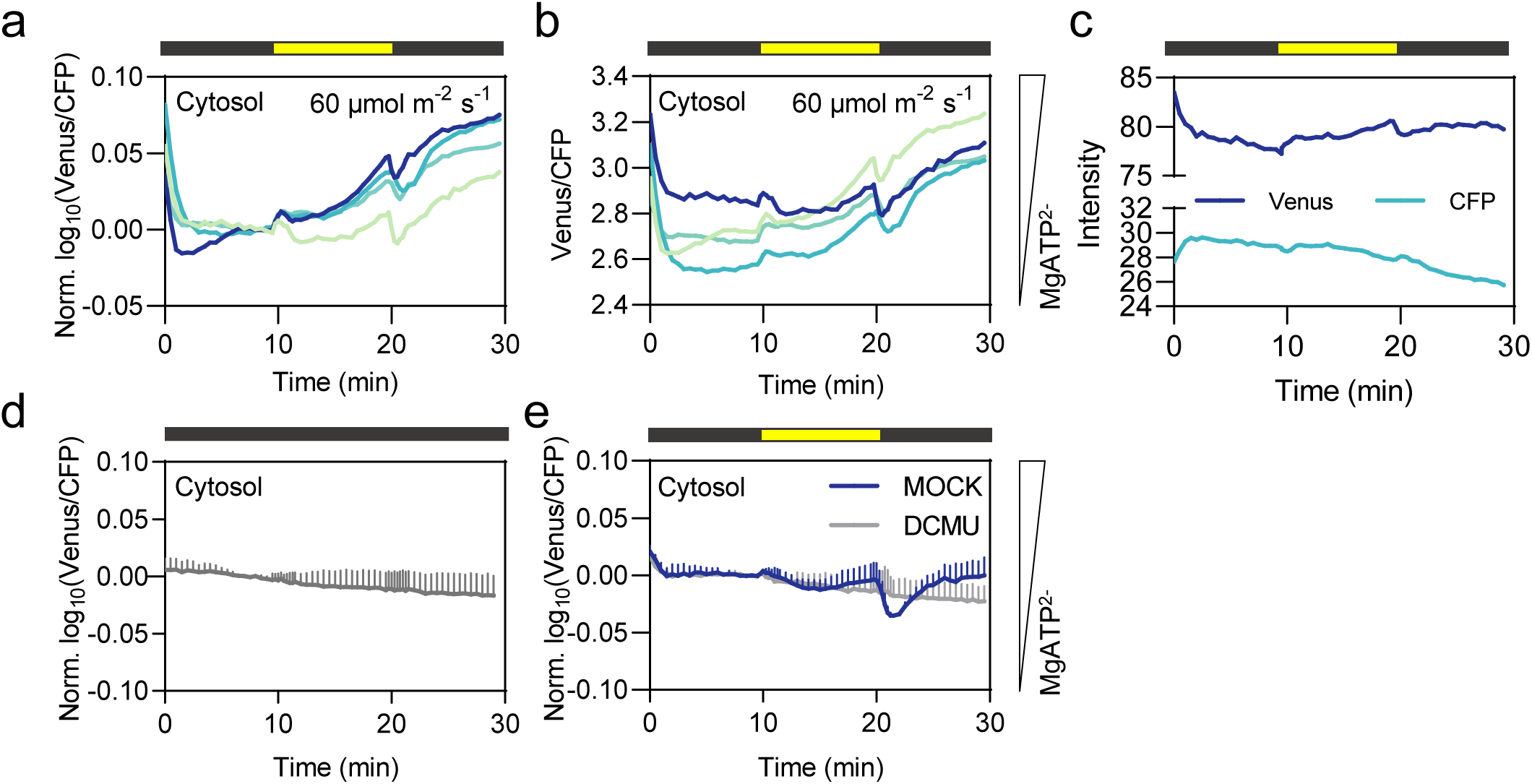
Single replicates and individual channels of cytosol-located ATeam1.03-nD/nA ratio under dark-light transitions (60 µmol m^-2^ s^-1^). All replicates showed cytosolic MgATP^2-^ increasing from dark to light transition and after illumination. In **(a)**, data are normalized to the average of the last five data points prior illumination and log10-transformed. In **(b)**, data are not normalized or log10- transformed. Different colors indicate single replicates. **c** Emission of single channels of cytosol-located ATeam1.03-nD/nA in response to dark-light transitions. n = 4. **d** ‘No light’ control to cytosol-located ATeam1.03-nD/nA. n = 7. **e** ’DCMU’ control of cytosol-located ATeam1.03-nD/nA: Leaf discs were pre-incubated in 20 µM DCMU and 0.2% (v/v) EtOH, or 0.2% (v/v) EtOH, respectively, for 35 to 45 min. Light intensity was 60 µmol m^-2^ s^-1^. n = 3. The black and yellow bars show the dark and light phases separately. Error bars = SD.

**Supplemental Figure 15.**
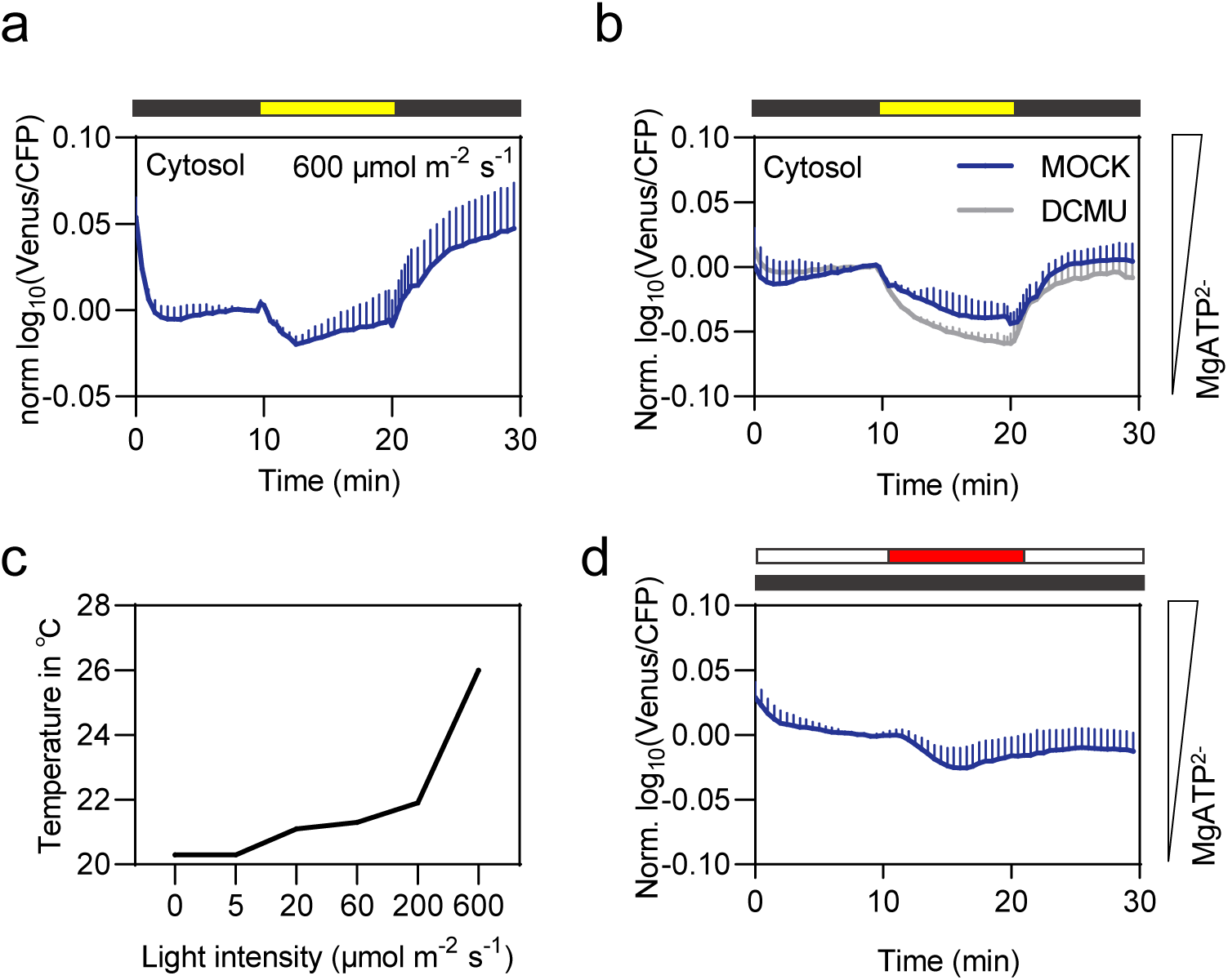
Cytosol-located ATeam1.03-nD/nA response to 600 µmol m^-2^ s^-1^. **a** Cytosol-located ATeam1.03-nD/nA line in response to 600 µmol m^-2^ s^-1^ light (indicated by yellow bars) for 10 min. n = 5. **b** ’DCMU’ control of cytosol-located ATeam1.03-nD/nA: Leaf discs were pre-incubated in 20 µM DCMU and 0.2% (v/v) EtOH, or 0.2% (v/v) EtOH, respectively, for 35 to 45 min. n = 3. Light intensity is 600 µmol m^-2^ s^-1^. Data are normalized to the average of the last five data points prior illumination, log10-transformed and averaged. **c** The temperature measured in the location of the disc samples in the on-stage illumination system under different light intensities. **d** Cytosol-located ATeam1.03-nD/nA ratio in response to increased temperature. The dark phase is indicated by a black bar, the 23 ℃ phase is indicated by the white section of the bars, the 26 ℃ phase is indicated by a red section of bar. N = 4. Data are normalized to the average of the last five data points of 23 ℃ phase, log10-transformed and averaged. Error bars = SD.

**Supplemental Figure 16.**
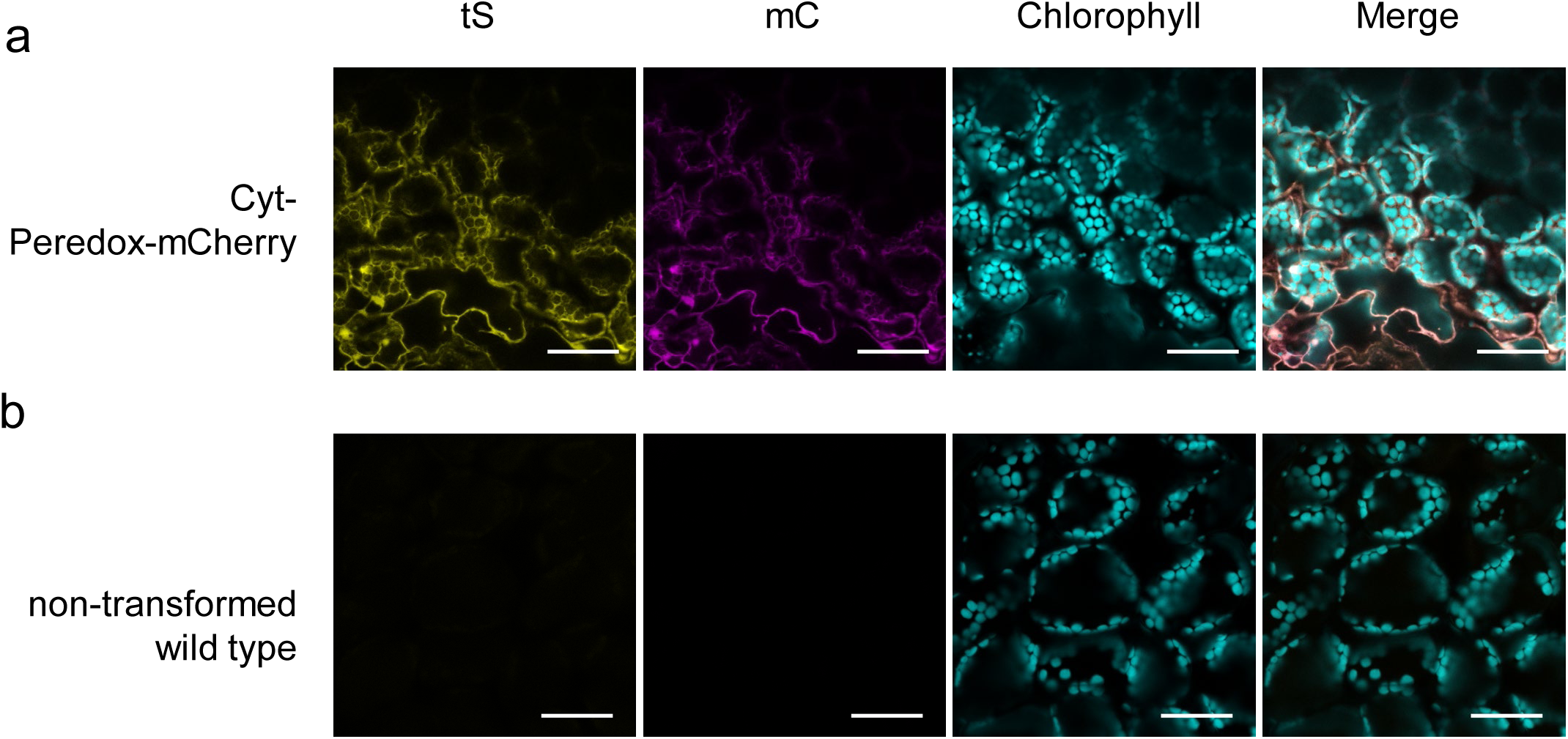
Confocal microscopy images of subcellular localization of Peredox-mCherry in the cytosol. **a** Confocal microscopy images of subcellular localization of Peredox-mCherry in the cytosol of cotyledon cells of 2-week-old Arabidopsis seedlings. **b** Non-transformed wild type imaged with the same confocal settings as in **(a)**, obtained from leaf discs of 4 to 5-week-old Arabidopsis plants. Confocal setting: excitation tS: 405 nm; emission tS: 499-544 nm; emission chlorophyll: 651-700 nm, excitation mC: 561 nm; emission mC: 588-624 nm. Scale bars: 50 µm. tS: tSapphire; mC: mCherry.

**Supplemental Figure 17.**
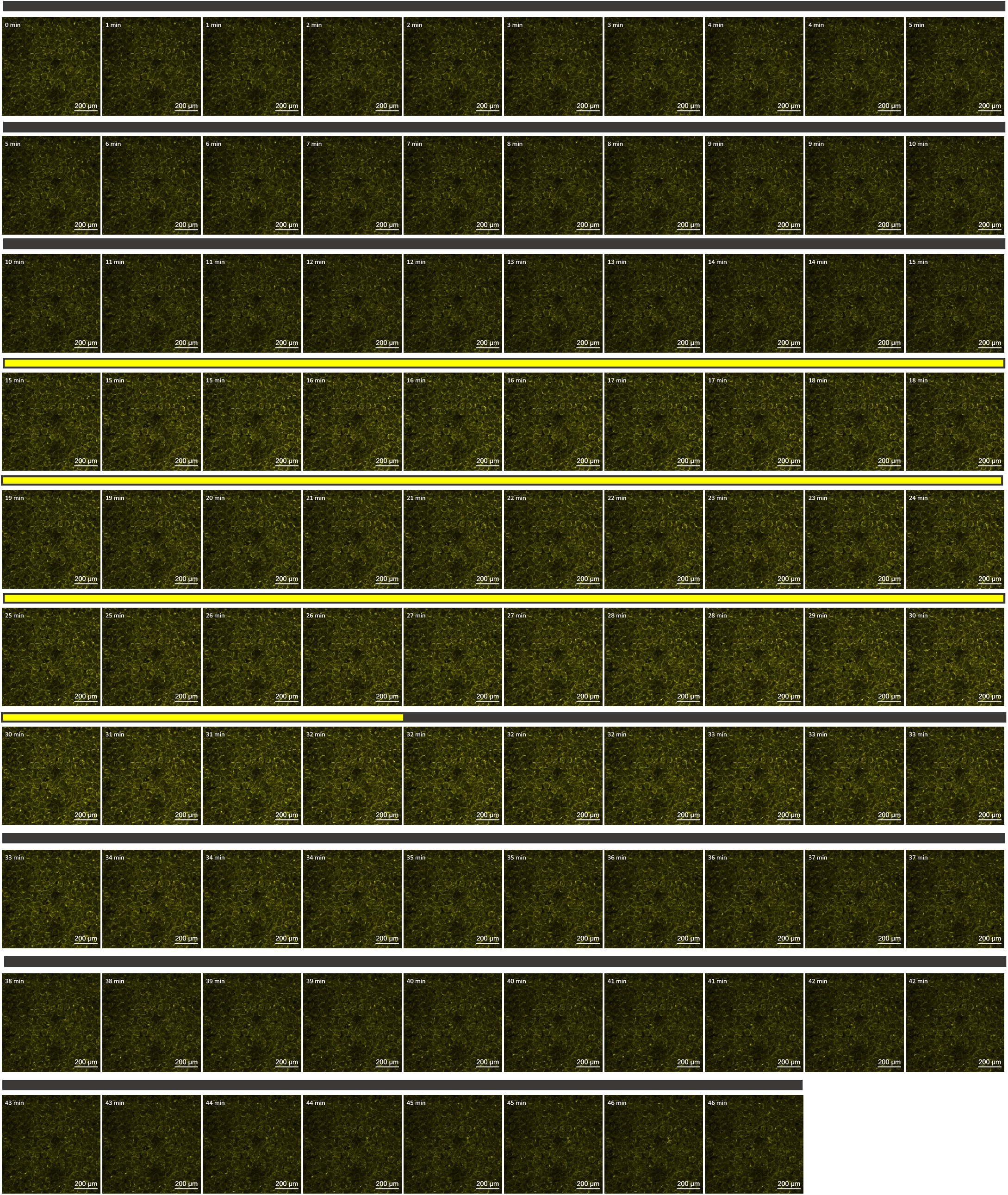
Confocal microscopy images of cytosol-located Peredox-mCherry in dark-light transitions. Confocal images of tSapphire (tS) channel of cytosol-located Peredox-mCherry at 405 nm excitation and 499-544 nm emission under 20 µmol m^-2^ s^-1^ dark-light transitions. The intensity of the yellow color indicated the extent of NAD reduction in the cytosol. The black and yellow bars show the dark and light phases separately.

**Supplemental Figure 18.**
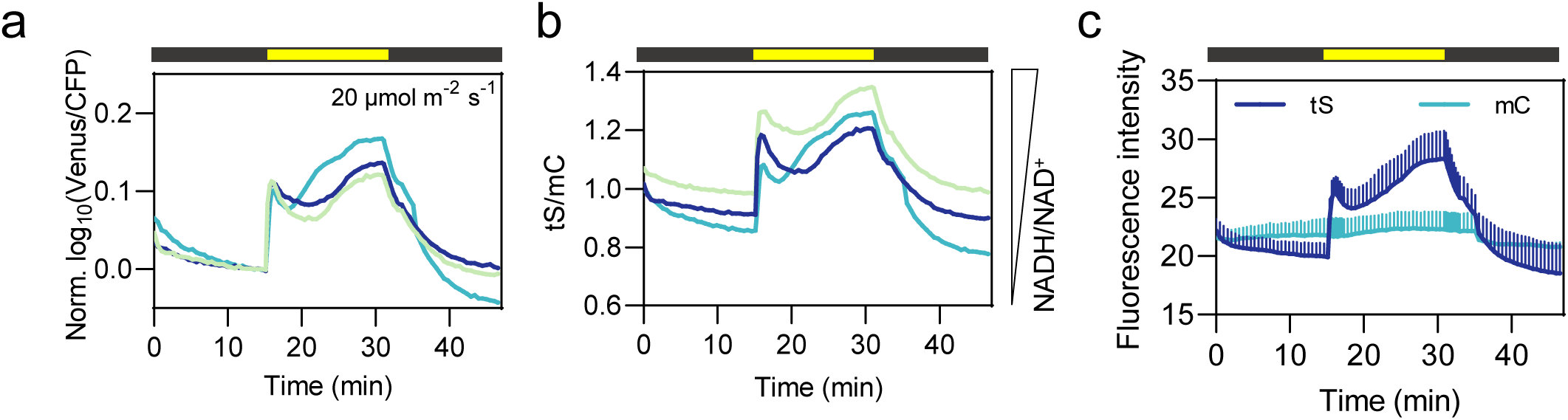
Single replicate and two separate channels of cytosol-located Peredox-mCherry under dark-light transitions (20 µmol m^-2^ s^-1^). **a, b** Three replicates of cytosol-located Peredox-mCherry showed similar dynamics under 20 µmol m^-2^ s^-1^ dark-light transitions. In **(a)**, data are normalized to the average of the last five data points prior illumination and log10-transformed. In **(b)**, data are not normalized or log10-transformed. **c** Emission of single channels of Peredox-mCherry in response to dark-light transitions. The black and yellow bars show the dark and light phases separately. Error bars = SD. tS: tSapphire; mC: mCherry.

**Supplemental Figure 19.**
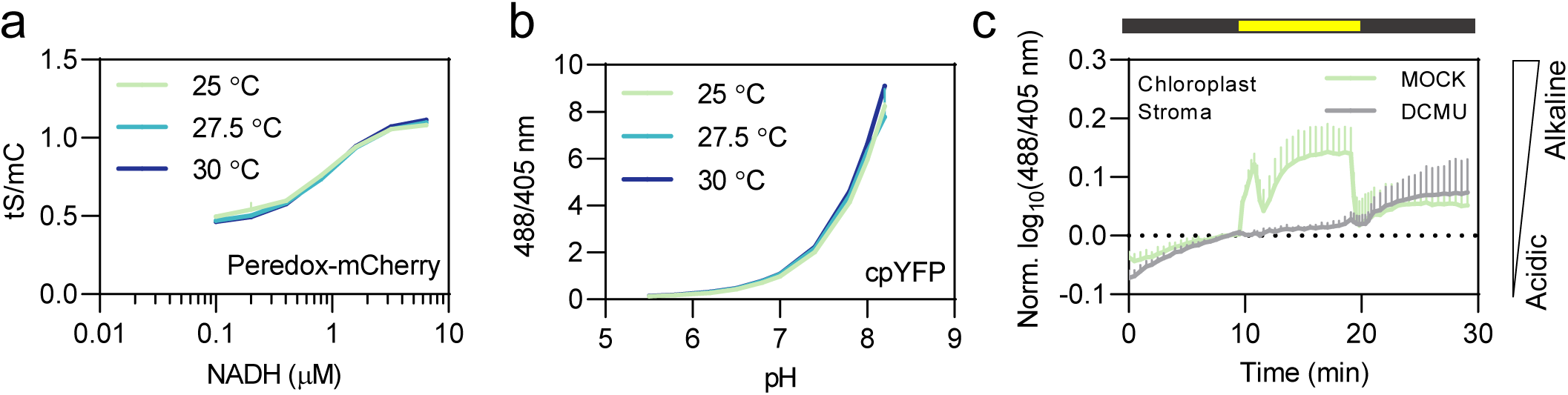
Temperature effect on NAD redox and pH sensor. *In vitro* assay of the temperature effect on Peredox-mCherry and cpYFP are done by Microplate Reader (BMG LABTECH, Ortenberg, Germany) and shown separately in **(a)** and **(b)**. In **(a)**, NADH titration is in the presence of 0.5 mM NAD^+^. All the samples are preincubated at corresponding temperatures at least 30 minutes before measurement, n=3. In **(c)**, ’DCMU’ control of chloroplast stroma located cpYFP: Leaf discs were pre-incubated in 20 µM DCMU and 0.2% (v/v) EtOH, or 0.2% (v/v) EtOH, respectively, for 35 to 45 min. n = 3. Light intensity was 600 µmol m^-2^ s^-1^, which has increased temperature. Data are normalized to the average of the last five data points in the first dark phase, log10-transformed and averaged. Error bars = SD.

## Supplemental Material

**Supplemental Table 1.**
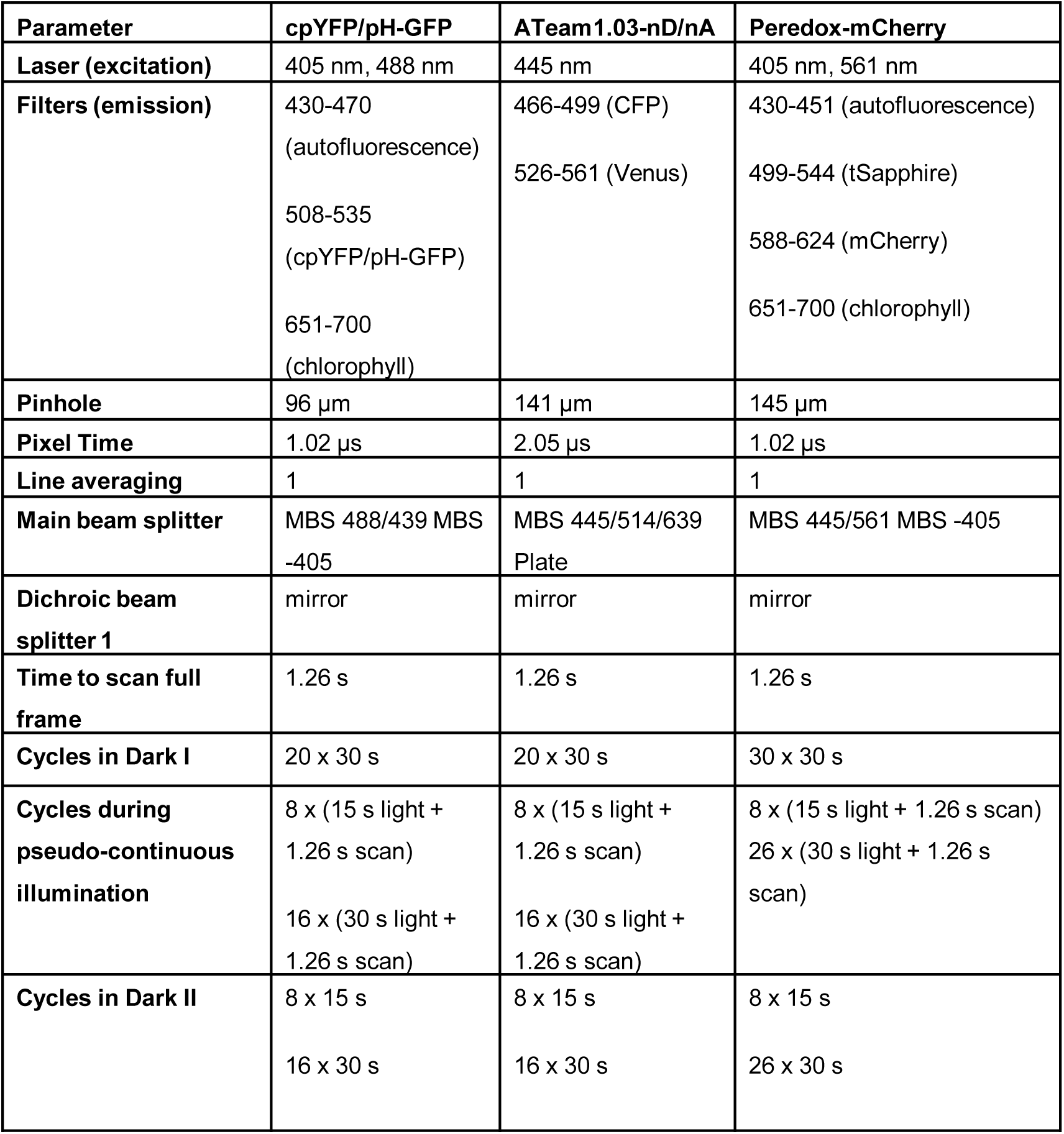
Microscopy parameters used for confocal imaging of cpYFP, pH-GFP, ATeam1.03-nD/nA and Peredox-mCherry in leaf mesophyll. Upper part of the table lists the parameters of the technical microscopic setup. The lower part gives details on the illumination regime applied for to the respective sensor.

| **Parameter** | **cpYFP/pH-GFP** | **ATeam1.03-nD/nA** | **Peredox-mCherry** |
| --- | --- | --- | --- |
| **Laser (excitation)** | 405 nm, 488 nm | 445 nm | 405 nm, 561 nm |
| **Filters (emission)** | 430-470<br>(autofluorescence)<br>508-535<br>(cpYFP/pH-GFP)<br>651-700<br>(chlorophyll) | 466-499 (CFP)<br>526-561 (Venus) | 430-451 (autofluorescence)<br>499-544 (tSapphire)<br>588-624 (mCherry)<br>651-700 (chlorophyll) |
| **Pinhole** | 96 $\mu\text{m}$ | 141 $\mu\text{m}$ | 145 $\mu\text{m}$ |
| **Pixel Time** | 1.02 $\mu\text{s}$ | 2.05 $\mu\text{s}$ | 1.02 $\mu\text{s}$ |
| **Line averaging** | 1 | 1 | 1 |
| **Main beam splitter** | MBS 488/439 MBS<br>-405 | MBS 445/514/639<br>Plate | MBS 445/561 MBS -405 |
| **Dichroic beam**<br>**splitter 1** | mirror | mirror | mirror |
| **Time to scan full**<br>**frame** | 1.26 s | 1.26 s | 1.26 s |
| **Cycles in Dark I** | 20 x 30 s | 20 x 30 s | 30 x 30 s |
| **Cycles during**<br>**pseudo-continuous**<br>**illumination** | 8 x (15 s light +<br>1.26 s scan)<br>16 x (30 s light +<br>1.26 s scan) | 8 x (15 s light +<br>1.26 s scan)<br>16 x (30 s light +<br>1.26 s scan) | 8 x (15 s light + 1.26 s scan)<br>26 x (30 s light + 1.26 s<br>scan) |
| **Cycles in Dark II** | 8 x 15 s<br>16 x 30 s | 8 x 15 s<br>16 x 30 s | 8 x 15 s<br>26 x 30 s |

Supplemental Video 1. Chloroplast stroma pH dynamics under dark-light transitions (60 µmol m^-2^ s^-1^) revealed by pH sensor cpYFP.

Ratiometric video of mesophyll expressing cpYFP in the chloroplast stroma under dark-light transitions (60 µmol m^-2^ s^-1^). 488/405 nm excitation ratio at 508-535 nm emission is indicated in false color. Scale bars: 200 µm (bottom right corner). Color scale is on the bottom left corner. Time scale is on the top left corner; 10 fps. Dark and light phases were shown on the top right corner.

Supplemental Video 2. Chloroplast stroma pH dynamics under dark-light transitions (600 µmol m^-2^ s^-1^) revealed by pH sensor cpYFP.

Ratiometric video of mesophyll expressing cpYFP in the chloroplast stroma under dark-light transitions (600 µmol m^-2^ s^-1^). 488/405 nm excitation ratio at 508-535 nm emission is indicated in false color. Scale bars: 200 µm (bottom right corner). Color scale is on the bottom left corner. Time scale is on the top left corner; 10 fps. Dark and light phases were shown on the top right corner.

Supplemental Video 3. Cytosolic pH dynamics under dark-light transitions (200 µmol m^-2^ s^-1^) revealed by pH sensor cpYFP.

Ratiometric video of mesophyll expressing cpYFP in the cytosol under dark-light transitions (200 µmol m^-2^ s^-1^). 488/405 nm excitation ratio at 508-535 nm emission is indicated in false color. Scale bars: 200 µm (bottom right corner). Color scale is on the bottom left corner. Time scale is on the top left corner; 10 fps. Dark and light phases were shown on the top right corner.

Supplemental Video 4. Cytosolic pH dynamics under dark-light transitions (600 µmol m^-2^ s^-1^) revealed by pH sensor pH-GFP.

Ratiometric video of mesophyll expressing pH-GFP in the cytosol under dark-light transitions (600 µmol m^-2^ s^-1^). 405/488 nm excitation ratio at 508-535 nm emission is indicated in false color. Scale bars: 200 µm (bottom right corner). Color scale is on the bottom left corner. Time scale is on the top left corner; 10 fps. Dark and light phases were shown on the top right corner.

Supplemental Video 5. Mitochondrial matrix pH dynamics under dark-light transitions (600 µmol m^-2^ s^-1^) revealed by pH sensor cpYFP.

Ratiometric video of mesophyll expressing cpYFP in the mitochondrial matrix under dark-light transitions (600 µmol m^-2^ s^-1^). 488/405 nm excitation ratio at 508-535 nm emission is indicated in false color. Scale bars: 200 µm (bottom right corner). Color scale is on the bottom left corner. Time scale is on the top left corner; 10 fps. Dark and light phases were shown on the top right corner.

Supplemental Video 6. Chloroplast stroma MgATP^2-^ dynamics under dark-light transitions (60 µmol m^-2^ s^-1^) revealed by ATP sensor ATeam1.03-nD/nA.

Ratiometric video of mesophyll expressing ATeam1.03-nD/nA in the chloroplast stroma under dark-light transitions (60 µmol m^-2^ s^-1^). The ratio of Venus/CFP is indicated in false color (excitation: 445 nm, Venus emission: 526-561 nm; CFP emission: 464-499 nm). Scale bars: 200 µm (bottom right corner). Color scale is on the bottom left corner. Time scale is on the top left corner; 10 fps. Dark and light phases were shown on the top right corner.

Supplemental Video 7. Cytosolic MgATP^2-^ dynamics under dark-light transitions (60 µmol m^-2^ s^-1^) revealed by ATP sensor ATeam1.03-nD/nA.

Ratiometric video of mesophyll expressing ATeam1.03-nD/nA in the cytosol under dark-light transitions (60 µmol m^-2^ s^-1^). The ratio of Venus/CFP is indicated in false color (excitation: 445 nm, Venus emission: 526-561 nm; CFP emission: 464-499 nm). Scale bars: 200 µm (bottom right corner). Color scale is on the bottom left corner. Time scale is on the top left corner; 10 fps. Dark and light phases were shown on the top right corner.

Supplemental Video 8. Cytosolic NADH/NAD^+^ dynamics under dark-light transitions (20 µmol m^-2^ s^-1^) revealed by NADH/NAD^+^ sensor Peredox-mCherry.

Ratiometric video of mesophyll expressing Peredox-mCherry in the cytosol under dark-light transitions (20 µmol m^-2^ s^-1^). The ratio of tSapphire/mCherry (tS/mC) is indicated in false color (tS: excitation: 405 nm, emission: 499-544 nm; mC: excitation: 561 nm, emission: 588-624 nm). Scale bars: 200 µm (bottom right corner). Color scale is on the bottom left corner. Time scale is on the top left corner; 10 fps. Dark and light phases were shown on the top right corner.

